# Mitochondrial cardiolipin sequestration of caspofungin underlies *Cryptococcus neoformans* inherent resistance and may contribute to cardiotoxicity

**DOI:** 10.64898/2026.01.20.700633

**Authors:** Anna K. Loksztejn, Rajendra Upadhya, Eduardo A. Caro, David M. Gooden, Adi Yona, Porter K. Ellis, Micha Fridman, Maria A. Schumacher, Richard G. Brennan, Maureen J. Donlin, Jennifer K. Lodge

## Abstract

Cryptococcus exhibits inherent resistance to the echinocandin, caspofungin, which inhibits the synthesis of (1,3)-β-D-glucan, a key component of the polysaccharide cell wall. The essential *FKS1* gene encodes the catalytic subunit of (1,3)-β-D-glucan synthase and caspofungin effectively inhibits its activity *in vitro,* yet the drug remains ineffective against Cryptococcus, suggesting mechanisms beyond target insensitivity. The underlying mechanisms of caspofungin resistance remain unknown, although altered regulation of cell-wall remodeling genes, plasma membrane modifications, drug efflux pathways, and melanin biosynthesis have been suggested. Using boron dipyrromethene (BD-) and fluorescein (F-) labelled caspofungin, we demonstrate that caspofungin enters the cryptococcal cell and primarily accumulates in the mitochondrial inner membrane rather than the plasma membrane. We further establish that this mitochondrial accumulation is driven by a specific interaction between caspofungin and cardiolipin, a phospholipid found in mitochondrial membranes. We demonstrate that this unforeseen localization indicates that mitochondrial sequestration diminishes the effective drug concentrations at the intended target. Notably, the interaction between caspofungin and cardiolipin also takes place in human cells, establishing a mechanistic connection to caspofungin-related cardiotoxicity. Our findings reveal a previously unrecognized mechanism of echinocandin resistance in Cryptococcus and emphasize cardiolipin as an important factor in caspofungin effectiveness.

## Introduction

Invasive fungal infections are a growing global health threat, particularly among immunocompromised populations, due to the limited number of safe and effective antifungal drugs. The opportunistic pathogen *Cryptococcus neoformans* is a leading cause of meningoencephalitis and accounts for ∼100,000 deaths annually worldwide^1^. Despite its clinical significance, no vaccine exists for cryptococcosis, and current antifungal therapeutical strategies are insufficient^2^.

The fungal cell wall is a dynamic structure absent in mammalian cells, which makes it an attractive target for antifungal development. The (1,3)-β-D-glucan synthase (GS), which catalyzes the synthesis of the important cell wall polysaccharide (1,3)-β-D-glucan, is the proven target of echinocandins^3,4^. GS is composed of the Fks1 catalytic subunit (FK506-supersensitive) and a regulatory subunit, Rho1. The existing glucan synthase-targeting antifungals are thought to inhibit glucan synthase activity by binding to the extracellular surface of the Fks1 subunit, however, the detailed mechanism by which these glucan synthase-targeting antifungals perturb glucan synthase function in cells is not fully understood^5^. Intriguingly, *C. neoformans* is intrinsically resistant to this drug class, despite the essentiality of its glucan synthase catalytic subunit^6,7^. Interestingly, prior studies have shown that partially purified cryptococcal glucan synthase can be inhibited by the echinocandin, caspofungin (CSF)^8^. Moreover, the viability of cryptococcal cells is impacted in the presence of CSF concentrations exceeding those achievable in patients, suggesting that glucan synthase remains a druggable target, provided that the CSF sensitivity can be increased. Several groups have performed extensive genetic screens to understand the mechanisms behind caspofungin tolerance. Among *C. neoformans* mutants identified, enhanced caspofungin efficacy was attributed to disruption of stress-response pathways^9^ as well as changes in membrane permeability, but not the expected defects in the cell wall^10^. Additionally, it was shown that interruption of lipid trafficking pathways sensitizes *C. neoformans* to caspofungin^11^ and that this mechanism can potentially be used to increase CSF efficacy by the introduction of appropriate drug modifications^12^. Under stress, cryptococcal cells have been shown to undergo melanization, which is a fungal defense mechanism involving the production of dark pigments that reduce drug, e.g. caspofungin, effectiveness. However, even melanized cells become susceptible to high caspofungin concentrations, suggesting that the CSF resistance mechanism is functional regardless of the melanization status of the cells^13^. Unmelanized *C. neoformans* is still ∼100-fold more resistant than susceptible *Candida* or *Aspergillus* sp. in MIC/MEC analysis using Clinical and Laboratory Standards Institute (CLSI) methods^8,14,15^. Combined, these experimental data point to highly complex mechanisms that are responsible for caspofungin tolerance in *C. neoformans*.

Efforts to develop novel glucan synthase inhibitors have been hindered by limited insight into drug–target interactions and critically, an incomplete understanding of cryptococcal resistance pathways. Derivatization of drugs with fluorescent probes is a useful approach in studying complex drug interactomes including the response of different fungi to echinocandins. Fluorescent derivatives of echinocandins have been developed to probe drug susceptibility and resistance in *Candida albicans* cells revealing that internalization of echinocandins into the lysosome leads to their reduced concentrations in the plasma membrane, which encompass the targeted β-(1,3)-D-glucan synthases, resulting in diminished antifungal activity^16,17^. A similar effect was observed for *Aspergillus fumigatus* probed with the same fluorescent caspofungin. Initially the probe accumulates in the interface between the cell membrane and cell wall and then internalizes into vacuoles^18^.

In the present study, we used fluorescently labelled caspofungin derivatives (boron dipyrromethene (BD-) and fluorescein (F-) conjugated) to probe drug localization in *C. neoformans* cells. Unexpectedly, we found that caspofungin accumulated predominantly in the mitochondrial membrane rather than the plasma membrane in which glucan synthase resides. These findings suggest that mitochondrial sequestration contributes to the reduced antifungal efficacy of caspofungin in *C. neoformans* and implicate mitochondrial stress responses in modulating drug susceptibility.

## Results

### Development of fluorescent BD-CSF probe

Fluorescently labelled probes are effective tools to probe the localization of molecules within a cell, particularly if the probe retains the biological activity of the underivatized molecule^19^. Previously, caspofungin (CSF) has been derivatized by attachment of the fluorescent probe to one of the primary amine groups (Fig. 1A, red) or attachment to the homo-tyrosine site, the chemical modification of which was reported not to significantly affect the drug efficacy (Fig. 1A, green)^11,16,17,20^. BODIPY-CSF obtained by amine group derivatization with boron dipyrromethene dye (BODIPY-) was used to characterize caspofungin uptake by *C. albicans*^20^, as well as to determine the role of a lipid flippase in *C. neoformans* virulence^11^. However, the previously described green BODIPY-CSF derivative^11,20^ was barely detectable in wild-type *C. neoformans* partially due to its low photostability^16^.

**Figure 1:**
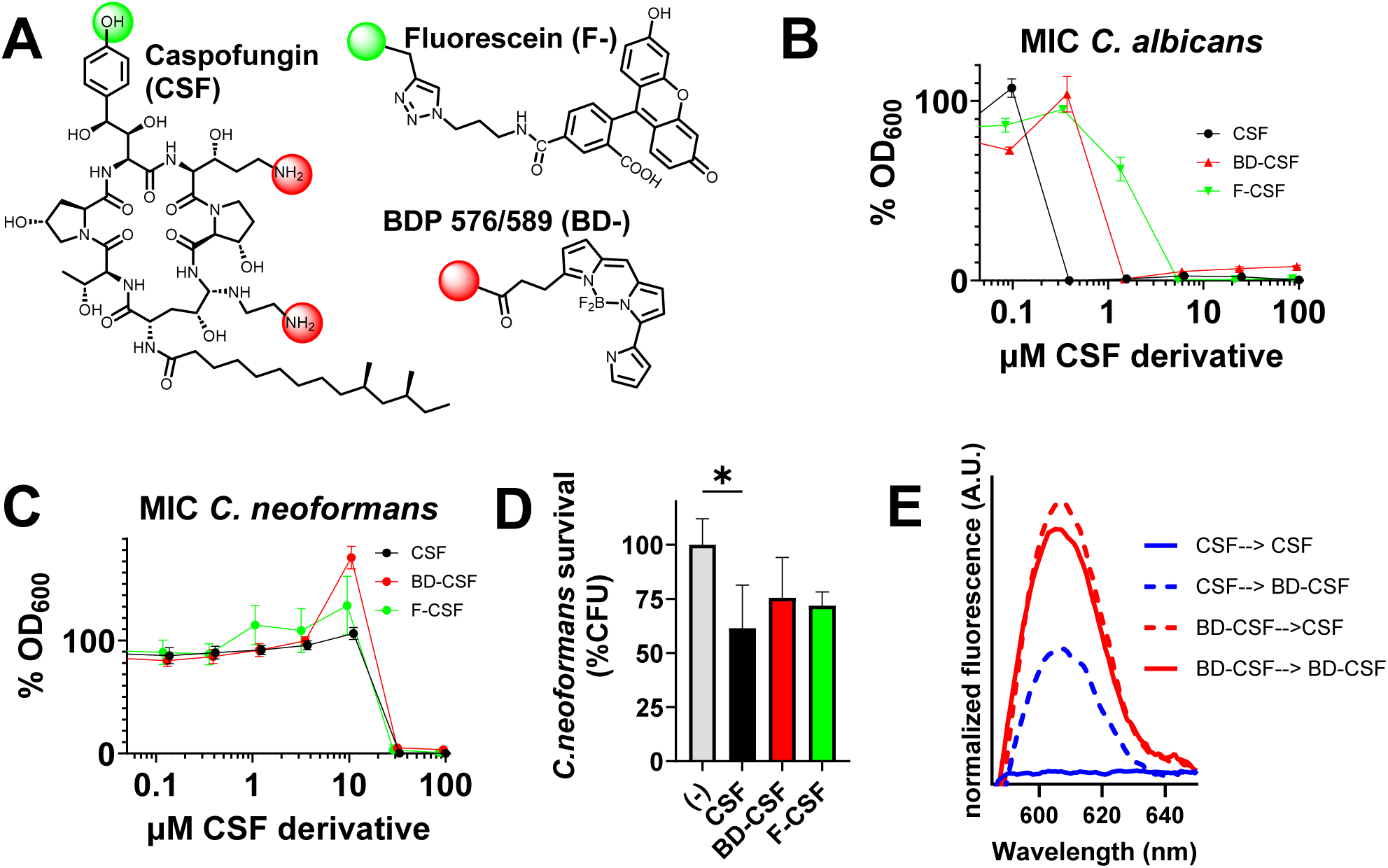
**A:** Chemical structure of caspofungin with marked positions utilized for derivatization with fluorescein (F-) and BDP 576/589 (BD-); amine groups (red); homo tyrosine (green). **B:** Determination of MIC for *C. albicans* **C**: and for *C. neoformans* **D**: Cell survival assay performed for cells treated with caspofungin (black) and its fluorescent derivatives – BD-CSF (red) and F-CSF (green). **E**: Spectral characterization of BD-CSF fluorescence bound to Cryptococcus cells following sequential treatment with CSF (blue) or BD-CSF (red), followed by either the same derivative (solid line) or the alternate derivative (dashed line).

We initiated our experiments using a BODIPY-CSF derivative. However, due to its low efficacy against *C. albicans*, complete lack of activity against *C. neoformans*, and the previously reported poor photostability of the BODIPY-CSF probe, we discontinued its use in subsequent experiments. Instead, we applied the same synthetic strategy involving amine group modification, but derivatized caspofungin with a different boron dipyrromethene analogue, a fluorophore that was potentially more photostable. Our design aimed to exploit the advantages of an alternative chemical structure, yielding a probe with high quantum yield and a long excited-state lifetime: the red-fluorescent BD-CSF (Fig. 1A). To generate this probe, caspofungin was derivatized at one of its primary amino groups (Supplementary Fig. 1A) using the N-hydroxysuccinimide ester of BDP® 576/589 (BD-NHS). The presence of mono-labelled BD-CSF was confirmed by LC–MS; however, we did not determine which of the two available amine groups in the caspofungin structure was derivatized (Supplementary Fig. 1B). Since the attachment of a hydrophobic fluorophore to one of the primary amines of caspofungin can potentially affect the chemical properties and efficacy of the compound, we also utilized an alternative synthetic approach and prepared the green fluorescent F-CSF derivative with the fluorescein (F-) label attached to the homo-tyrosine site (Fig. 1A, Supplementary Fig. 2) according to the previously published synthetic protocol^17^.

Because derivatization of caspofungin with fluorescent probes alters its chemical structure, we anticipated potential differences in antifungal efficacy between the fluorescent CSF derivatives and the parent compound, as previously reported for *C. albicans*^16^. To assess whether derivatization compromised activity, we determined the minimum inhibitory concentration (MIC) values for both fluorescent CSF derivatives and compared them with unlabeled caspofungin. The MIC was defined as the lowest concentration at which growth was completely inhibited relative to the no-drug control.

Control experiments were performed with the CSF-sensitive *C. albicans* WO1 strain (Fig. 1B) and compared with wild-type *C. neoformans* KN99 (Fig. 1C) using previously established assay conditions ^10,21^. The reduced efficacy of the fluorescent derivatives against *C. albicans* was consistent with earlier findings for analogous probes tested across multiple *C. albicans* strains (Fig. 1B)^16^. Although BD-CSF and F-CSF were significantly less effective in *C. albicans*, both derivatives exhibited comparable activity against *C. neoformans*. The MIC values obtained (32 µM for BD-CSF and 29 µM for F-CSF) were similar to that of the parent drug (33 µM) (Fig. 1C) and aligned with previously published MIC values ^8,13^.These inhibition assays reaffirm the inherent resistance of *C. neoformans* to caspofungin (approximately 100-fold greater than *C. albicans*) and demonstrate that MIC values exceed clinically relevant plasma concentrations (∼10 µM)^22^. Importantly, BD-CSF and F-CSF displayed comparable effects on cryptococcal cell density to caspofungin, thereby validating the successful design and development of these fluorescent caspofungin probes.

We next compared the fungicidal activity of caspofungin and its fluorescent derivatives using a fungal survival assay. Wild-type *C. neoformans* (KN99) cells were grown to mid-logarithmic phase and then treated with ∼MIC concentrations of CSF, BD-CSF, or F-CSF for 24 hours, after which cell survival was quantified by CFU enumeration. As shown in Fig. 1D, underivatized caspofungin resulted in ∼40% killing, whereas the CSF derivatives did not induce significant fungicidal activity, though both showed a modest downward trend, reaching approximately 30% reduction in viability. (Fig. 1D).

We also performed spectral characterization of the BD-CSF probe bound to cryptococcal cells following a subsequent treatment with fluorescent and then underivatized caspofungin (Fig. 1E). Notably, pretreatment with underivatized caspofungin reduced the amount of the BD-CSF fluorescence signal bound to cryptococcal cells, suggesting that CSF and BD-CSF are likely accumulating in the same location. However, sequential treatment of cells—first with BD-CSF and then with unmodified caspofungin produced a fluorescence signal comparable to the BD-CSF–only control, suggesting that the fluorescent derivative of caspofungin has a higher affinity to the drug binding site without increasing its efficacy.

Fluorescence microscopy of cells treated with caspofungin derivatives revealed that both CSF probes localized inside the yeast cells, rather than at the expected outer membrane site where the glucan synthase complex resides. (Fig. 2A). Moreover, we observed two prominent phenotypic features, dispersed and punctate fluorescence, within the cells, that appeared consistently across all samples, irrespective of which CSF derivative was used prior to microscopic analysis. Statistical distribution of these phenotypes revealed that a substantial fraction of the population (59% and 62% of cells for BD-CSF and F-CSF respectively) displayed dispersed BD-CSF/F-CSF fluorescence throughout the cell and often the cells have an inner ring of fluorescence as well as a fainter outer ring, which may be associated with the plasma membrane. In contrast, approximately 40% of cells exhibited a punctate fluorescence pattern, resembling stress granules^23^ or distinct mitochondrial phenotypes^24^ previously observed in cryptococcal cells. To assess the proximity of the fluorescent caspofungin signal to the cell wall, we co-stained cells with calcofluor white (CFW), which binds to cell wall chitin. The results confirmed that the probes are not primarily localized at the plasma membrane (Fig. 2A). To verify that the observed intracellular fluorescent signals did not originate from the nascent BD-NHS or BD-fluorescent moiety, we treated cryptococcal cells with BD-NHS and glycine derivatized with BD-NHS (BD-Gly), simulating the BD-CSF probe, for 24 hours at a concentration of 100 µM, which exceeds our standard experimental conditions. We did not find any uptake of the BD-NHS or BD-Gly probes. However, with BD-NHS, we observed staining within the cell wall, likely due to amine-group labelling of cell-wall proteins (Supplementary Fig. 3A).

**Figure 2:**
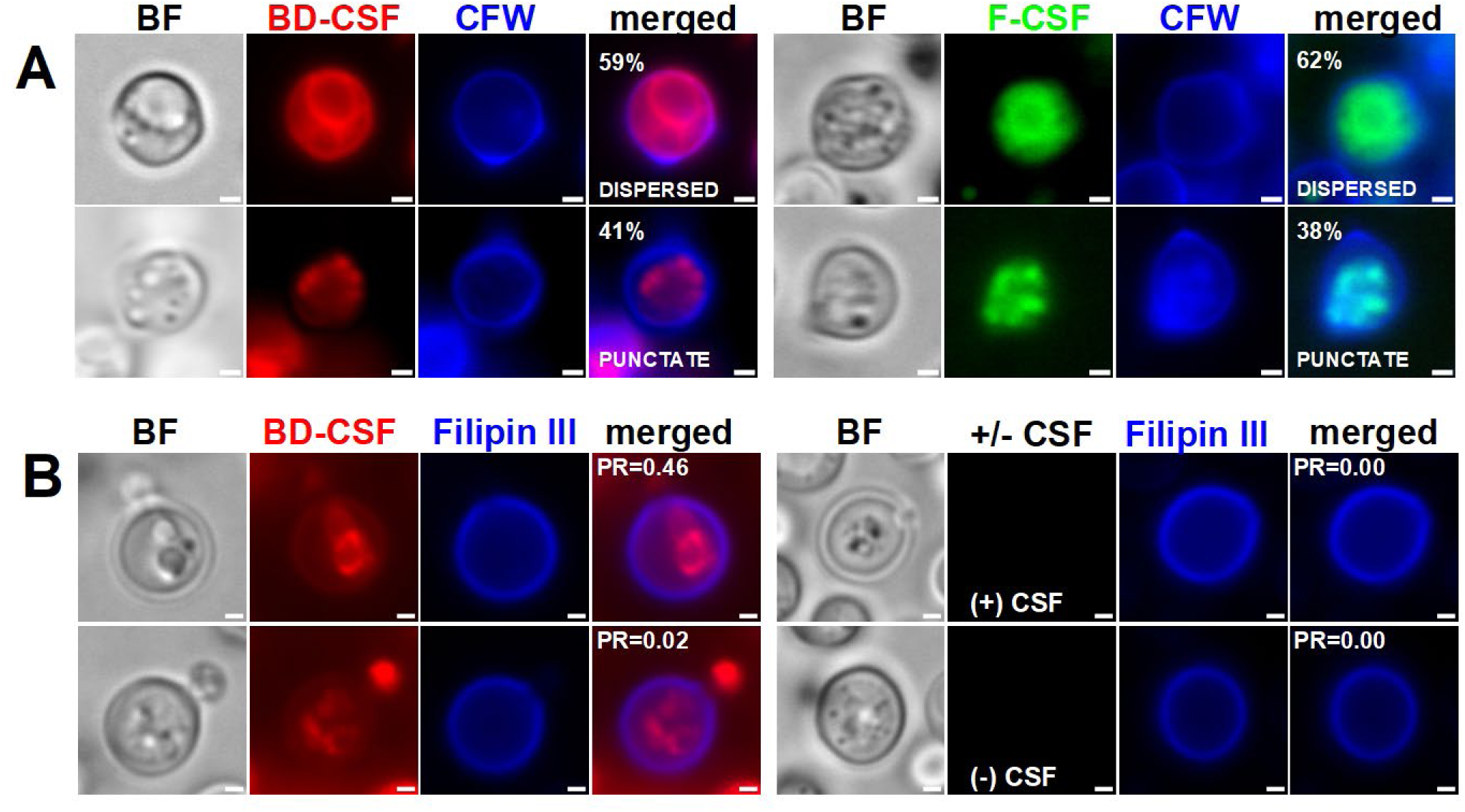
**A**: Representative images of wild-type *C. neoformans* cells treated with of BD-CSF (red) or F-CSF (green) exhibiting dispersed and punctate phenotypes with their respective abundance in the cell population. **B**: Wild-type *C. neoformans* cells stained with Filipin III (blue) following the treatment with BD-CSF (red) or +/- CSF as a control. Person’s coefficient values R (PR) describing colocalization of the Filipin III in the DAPI channel with corresponding fluorescence in the TRITC channel. Scale bar within each image marks 1µm length.

### BD-CSF fluorescence does not considerably overlap with the plasma membrane

The target of caspofungin, Fks1/glucan synthase, resides in the plasma membrane. The caspofungin structure contains a lipid tail that facilitates membrane binding^25^, proximal to the inhibition site. To determine whether BD-CSF co-localizes with the plasma membrane and to assess the impact of CSF on membrane integrity, we employed Filipin III, a sterol-binding fluorescent dye widely used as a reporter for plasma membrane structure and ergosterol content^26,27^ (Fig. 2B). Examination of a broad field of cells revealed heterogeneous staining (Supplementary Fig. 3B): some plasma membranes stained brightly with Filipin III, while others did not, even in the absence of CSF treatment. This variability is likely attributable to the low proportion of actively budding, metabolically active cells in saturated cultures grown in minimal media following prolonged exposure to caspofungin and BD-CSF.

No significant defects in outer membrane structure were observed in cells uniformly stained with Filipin III after treatment with CSF or BD-CSF (Fig. 2B). To quantify co-localization, we calculated Pearson’s correlation coefficient (R) for BD-CSF–treated cells. A coefficient of 1 indicates perfect co-localization of fluorescence signals, whereas 0 denotes no overlap. Only partial signal overlap was observed between BD-CSF and Filipin III (Fig. 2B), suggesting that the most of fluorescent caspofungin localizes to intracellular regions in those selected cells. Interestingly, this partial overlap was more prominent in the cells exhibiting the dispersed BD-CSF staining phenotype (Fig. 2B top left) as compared to the perceived punctate phenotype (Fig. 2B bottom left). Although these data are qualitative, due to uneven staining efficiency, these observations suggest that fluorescent CSF probes preferentially accumulate in internal structures, prompting further experiments to investigate the localization of the internalized probe.

### BD-CSF fluorescence colocalizes with MitoTracker dye

We observed that the fluorescence patterns of BD-CSF and F-CSF closely resembled the mitochondrial phenotypes - diffuse, fragmented, and tubular - in cells exposed to oxidative stress^24^. Because the membrane-anchoring lipid chain of caspofungin could facilitate its association not only with the plasma membrane but also with lipid-rich organelles such as mitochondria, we next examined if the fluorescent caspofungin probes localize to mitochondria. To this end, cryptococcal cells were incubated with CSF, BD-CSF, or F-CSF, followed by the carbocyanine-based dye MitoTracker, which enables mitochondrial visualization by fluorescence microscopy (Fig. 3A–B; Supplementary Fig. 4A–B). Interestingly, we reproduced MitoTracker derived mitochondrial phenotypes (diffuse, fragmented, tubular) as described previously by Chang *et al* ^24^ in cells pretreated with underivatized caspofungin as well as with the fluorescent probes (Fig. 3A-B; Supplementary Fig. 4A–B). Moreover, in cells pretreated with the fluorescent caspofungin analog BD-CSF followed by the MitoTracker staining, the dispersed fluorescence pattern derived from BD-CSF fluorescence aligned with the diffuse phenotype visualized by the MitoTracker dye, and the punctate fluorescence pattern of BD-CSF corresponded to the MitoTracker derived fragmented phenotype (Fig. 2A, Fig. 3B). Notably, the tubular phenotype we observed using MitoTracker did not have a corresponding BD-CSF derived fluorescence pattern. To quantify these mitochondrial phenotypic changes, we analyzed more than 100 individual cells treated with caspofungin or BD-CSF and compared the distribution of diffuse, fragmented and tubular phenotypes with that of untreated cells. Drug-treated cells displayed a markedly increased proportion of fragmented mitochondria, an indicator of heightened cellular stress (Fig. 3C–D). In addition, both caspofungin and BD-CSF–treated populations exhibited an increased number of budding cells, a finding which is elaborated below (Supplementary Fig. 4D).

**Figure 3.**
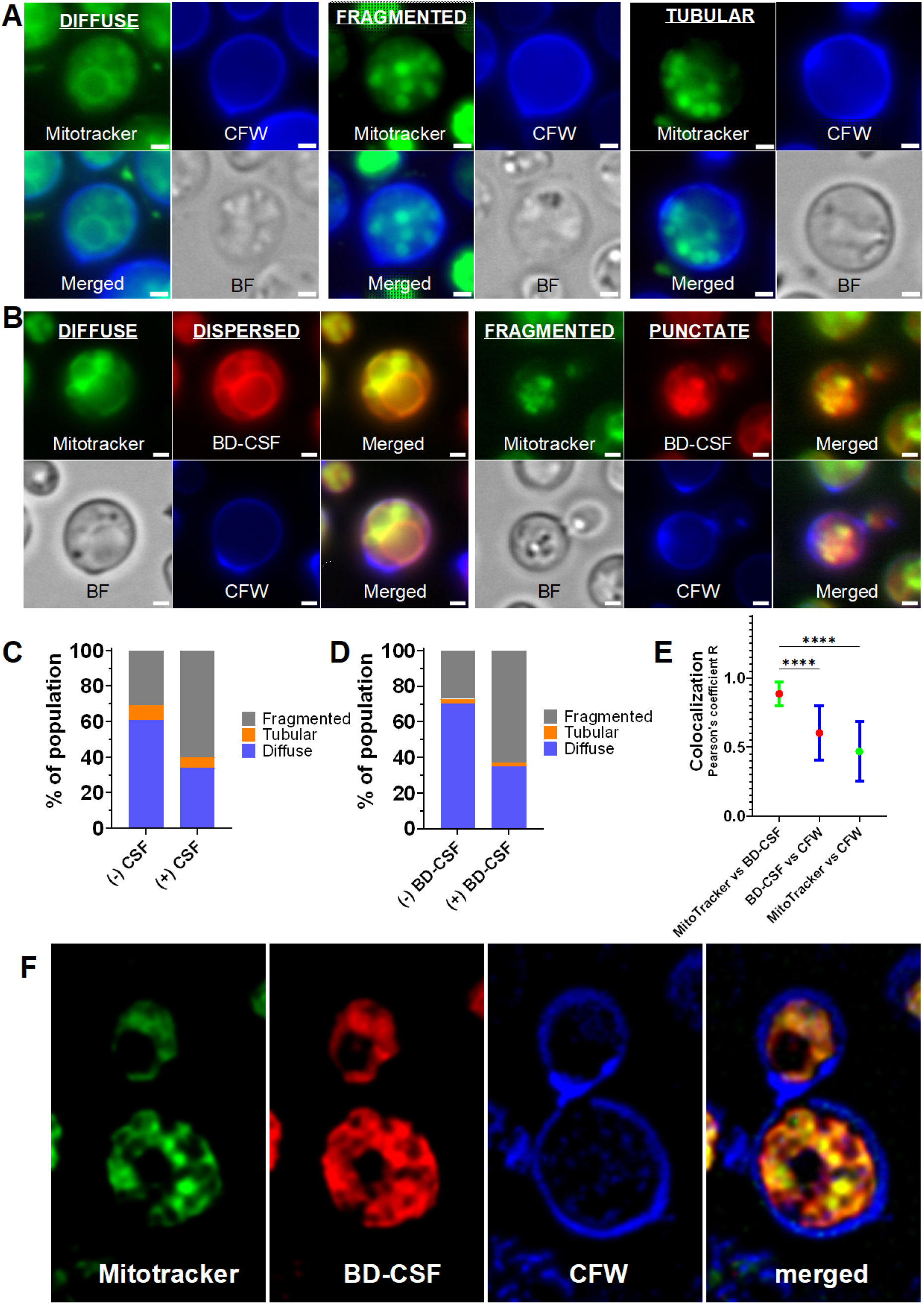
Representative images of wild-type *C. neoformans* cells treated with **A**: CSF and **B**:BD-CSF (red) co-labelled with MitoTracker (green) and calcofluor-white (blue) exhibiting different phenotypes present in the cell population. Scale bar within each image marks 1µm length. Phenotypic distribution of cells treated with **C:** CSF and **D:** BD-CSF. **E:** Average Pearson’s coefficient R calculated for colocalization of MitoTracker vs BD-CSF fluorescence (red marker, green error bars), BD-CSF vs CFW fluorescence (red marker, blue error bars), MitoTracker vs CFW fluorescence (red marker, blue error bars). **F:** Representative images of wild-type *C. neoformans* cells treated with BD-CSF (red) co-labelled with MitoTracker (green) and calcofluor-white (blue) and visualized using a high-resolution Elyra 7 microscope.

Closer examination of BD-CSF fluorescence revealed clear colocalization with MitoTracker staining (Fig. 3B); specifically, merged images produced a yellow signal indicating mitochondrial association of the fluorescent caspofungin probes. Given this overlap and the comparable phenotype distributions, we performed quantitative colocalization analysis (Fig. 3E). Pearson’s correlation coefficients (R) were calculated for MitoTracker and BD-CSF fluorescence from microscopy images of more than 100 individual cells per condition. As controls, we also computed Pearson’s coefficients for correlations of MitoTracker and BD-CSF each with CFW fluorescence. The average Pearson’s coefficient of ∼0.9 for BD-CSF and MitoTracker demonstrates strong colocalization (Fig. 3E). In contrast, significantly lower R values, ∼0.6, for BD-CSF–CFW overlap support only partial colocalization between plasma-membrane-associated BD-CSF fluorescence and CFW. The very low MitoTracker-CFW correlation, <0.5, is consistent with the spatial separation of mitochondria and the cell wall. To further validate mitochondrial colocalization of fluorescent caspofungin, we acquired high-resolution images of cells treated with BD-CSF and F-CSF (Fig. 3F; Supplementary Fig. 4C). Fluorescence from both probes strongly overlapped with MitoTracker, providing additional evidence that caspofungin localizes to mitochondria.

### Mitochondrial lipid reporter – Nonyl Acridine Orange- outcompetes BD-CSF fluorescence

We hypothesized that caspofungin, with a lipid side chain that facilitates membrane association, may be interacting with the lipids in the mitochondria. The phospholipid cardiolipin is located in the inner mitochondrial membrane. Cardiolipin binds to Nonyl Acridine Orange (NAO), a green fluorescent probe, in a 2:1 stoichiometric ratio^28^. NAO serves as a dependable marker that is independent of membrane potential. We conducted colocalization of BD-CSF with NAO to evaluate if the observed mitochondrial association is driven by lipids. Fluorescence microscopy revealed an apparent competition between NAO and BD-CSF, resulting in a partial reduction in the BD-CSF signal (Fig. 4A), indicating a potential interaction between CSF and cardiolipin. Analysis of images obtained prior to and following NAO staining of BD-CSF–treated cells revealed a statistically significant decrease in red fluorescence (Fig. 4B). Furthermore, the NAO signal was significantly diminished in BD-CSF–treated cells, (Fig. 4C) relative to untreated controls under the same conditions. This provides additional evidence for a direct interaction between caspofungin and cardiolipin.

**Figure 4.**
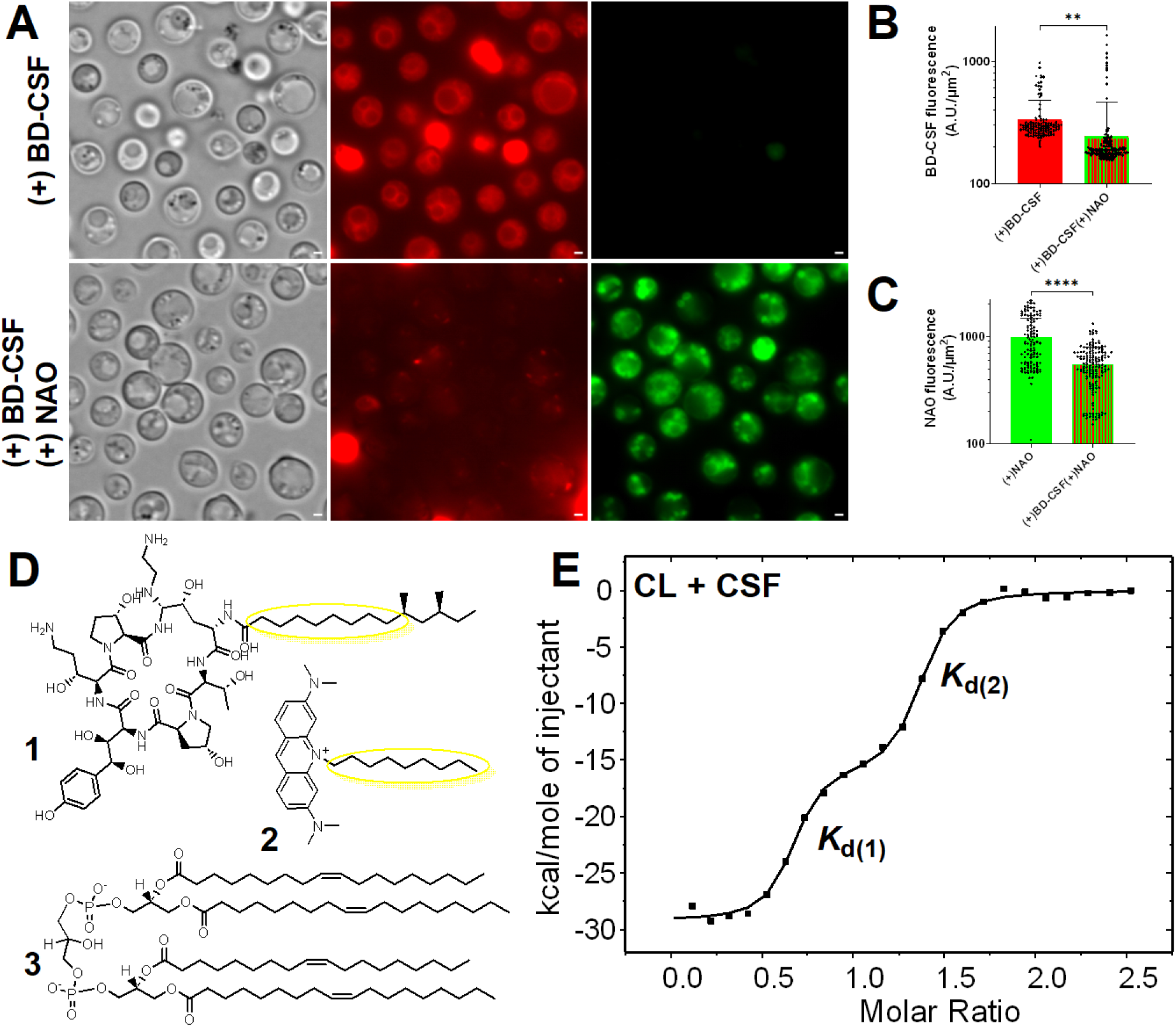
**A:** Representative wide field images of wild-type *C. neoformans* cells treated with BD-CSF (red) prior (top) and after (bottom) staining with NAO (green). Scale bar within each image marks 1µm length. **B:** Decrease in BD-CSF fluorescence following NAO treatment **C:** Decreased NAO uptake for cells pre-treated with BD-CSF in comparison with untreated cells **D:** Chemical structures of CSF (1) and NAO (2) both of which contain chemically similar lipid moieties. Chemical structure of 18:1 Cardiolipin (3) **E:** Representative isothermal titration calorimetry binding isotherm showing two-site binding between caspofungin (CSF) and cardiolipin (CL).

### Caspofungin binds to cardiolipin *in vitro*

Our imaging data indicating BD-CSF/NAO competition led us to hypothesize that cardiolipin may bind CSF. Since the interaction of NAO with cardiolipin has been previously documented ^28^, we compared the chemical structures of caspofungin and NAO in relation to cardiolipin and observed structural similarities in the lipid moieties of caspofungin and NAO that might be related to the cardiolipin-binding properties of these compounds. We reasoned this shared structural feature may underpin the shared cardiolipin-binding properties of these molecules (Fig. 4D). Additionally, it has been established that the length of the lipid chains is a significant factor influencing the properties of antifungal drugs, including caspofungin^29^. To test whether caspofungin binds to cardiolipin, we carried out a series of isothermal titration calorimetry (ITC) experiments to determine their *in vitro* interaction and the binding affinity and stoichiometry of the caspofungin-cardiolipin complex. ITC quantitatively assesses the binding thermodynamics of molecules and provides information on binding stoichiometry. Since the cardiolipin structure varies between different organisms, and the chemical structure of cryptococcal cardiolipin is unknown, we used the commercially available 18:1 cardiolipin which is the predominant form of cardiolipin in *S. cerevisiae* mitochondria^30^. The experiment was conducted in triplicate at 298K. In each experimental replicate, a 0.5 mL syringe containing a solution of cardiolipin (1.0 mM in EtOH/H_2_O = 1/1) was sequentially injected into a 2.50 mL sample cell containing a solution of caspofungin (25.0 μM in EtOH/H_2_O = 1/1) with stirring at 300 rpm. After data normalization, the equilibrium dissociation constants, *K*_d_, were calculated using a two-site model (Fig. 4E, Supplementary Fig. 5A). The average *K*_d_ values suggest a strong affinity of caspofungin for cardiolipin, with *K*_d(1)_ at 3.26 × 10^-9^ ± 5.54× 10^-10^ M and *K*_d(2)_ at 7.39 × 10^-7^ ± 4.93 × 10^-7^ M. Analogous to specific NAO-cardiolipin binding, the caspofungin-cardiolipin binding ratio is ∼1.5:1, consistent with a 2:1 stoichiometry observed for the NAO-cardiolipin interaction. The experimental deviations in the calculated stoichiometries likely arise from errors in ligand concentrations. Because cardiolipin is specific to mitochondria but is not its predominant phospholipid^30^, control experiments were conducted with phosphatidylcholine (PC; 1.0 mM, EtOH/H_2_O = 1/1) the predominant phospholipid in the mitochondrial membrane^30^ (Supplementary Fig. 5 B). The results obtained were inconsistent, and the considerably higher *K*_d_ values indicated a significantly weaker or nonexistent interaction that required a one-site model for curve fitting between caspofungin and phosphatidylcholine.

### Pretreatment of cells with a cardiolipin binding reagent enhances the efficacy of caspofungin

We hypothesized that the binding of caspofungin to cardiolipin in the mitochondrion results in a decreased effective concentration of the drug at the plasma membrane, where Fks1 is located. If this is the case, the result would be a diminished efficacy of caspofungin (Fig. 5A). Further, this would predict that this effect may be mitigated by pre-saturating available cardiolipin sites with a cardiolipin binding reagent, thereby preventing caspofungin from interacting with mitochondria while simultaneously increasing the effective concentration of caspofungin in the plasma membrane (Fig. 5B). We utilized NAO as proof of principle to achieve that effect. A competition assay was designed in which wild-type *C. neoformans* cells were initially cultured to an OD_600_ of 0.3 in YNB media at pH 7, followed by pre-incubation with 11 µM NAO per 10^8^ cells/mL. After a 1-hour incubation with NAO, cells were treated with 10 µM caspofungin or BD-CSF for 2 hours and 24 hours, followed by characterization using fluorescent microscopy. A significant decrease in mitochondrial BD-CSF uptake was observed in cells exposed to NAO compared to those not exposed (Fig. 5C). An increase in BD-CSF fluorescence was observed at the plasma membrane, likely colocalized with Fks1. Furthermore, following a three-hour exposure to NAO and caspofungin, there was a notable (89-fold) reduction in the viability of cells that had been pre-incubated with NAO and subsequently treated with caspofungin as compared to untreated control cultured simultaneously (Fig. 5D). The fluorescent caspofungin derivative, BD-CSF, reduced cell viability by a factor of 31 within a 3-hour period. The combined effects of NAO and caspofungin/BD-CSF treatments were more significant than those of NAO or caspofungin/BD-CSF administered individually. Given that prolonged, 24-hour incubation with NAO exhibits a cytostatic effect on cells (Supplementary Fig. 6A), we concentrated on the analysis of the data obtained from the earlier 3h timepoint. We examined alterations in the average NAO fluorescence signal subsequent to the caspofungin and BD-CSF treatments (Supplementary Fig. 6B). The average NAO fluorescence of cells remains largely unchanged following treatment with caspofungin. However, there is a significant decrease in the NAO fluorescence upon exposure to BD-CSF, indicating that BD-CSF is a stronger competitor for cardiolipin binding than caspofungin and can outcompete NAO from mitochondria. These data provide a likely explanation of reduced efficacy of BD-CSF during the treatment of NAO pre-saturated cells. We also analyzed the size changes of the budding and single cell populations under all experimental conditions (Supplementary Fig. 6C). A statistically significant reduction in cell size was noted in the budding cell population, with the most notable difference observed in cells pre-treated with NAO and subsequently exposed to caspofungin treatment (both CSF and BD-CSF). In single cells, only simultaneous treatment with NAO and BD-CSF lead to a statistically significant reduction in cell size. These results are consistent with the increased sensitivity of young, actively budding cells to antifungals^31-33^.

**Figure 5.**
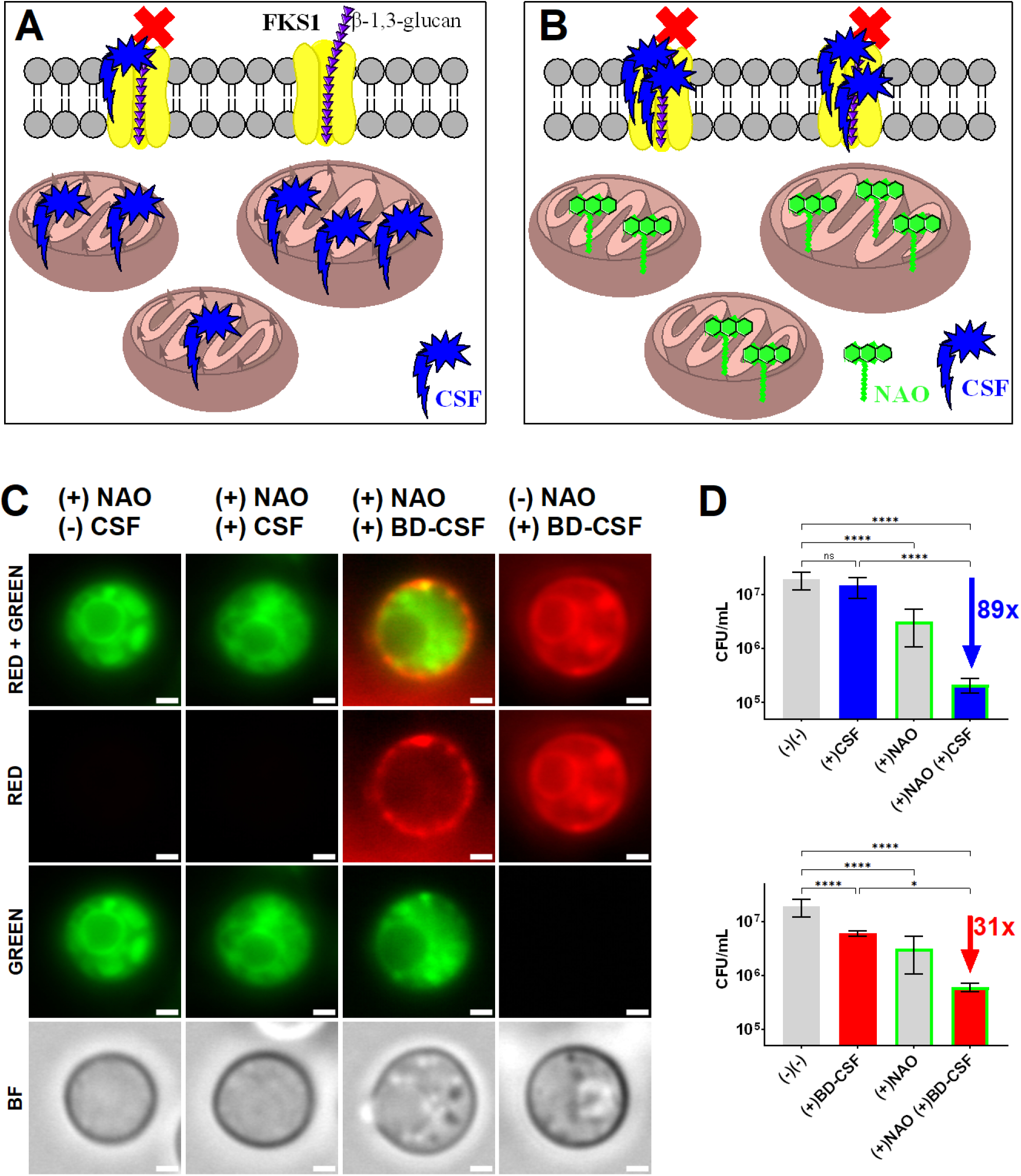
**A:** Schematic model of caspofungin distribution between its intended target - Fks1, and the off-site target mitochondrial cardiolipin implicating mitochondria are major sequesters of the available drug. **B:** Experimental design illustrating how pre-saturation of cardiolipin sites with the cardiolipin specific dye-NAO (green) increases the effective CSF concentration at its intended target, Fks1 **C:** Representative images of cells pre-treated with NAO prior to BD-CSF treatment showing increased plasma membrane localization of the drug. **D:** Viability assay of wild-type *C. neoformans* (gray) treated with CSF (blue) or BD-CSF (red), with or without NAO pre-treatment (green frame). After a cumulative 3h NAO pre-treatment followed by exposure to 10µM caspofungin (CSF) an 89-fold reduction in viability is observed, while NAO pre-treatment followed by 10 µM BD-CSF led to a 31-fold reduction, relative to the untreated control.

### Caspofungin binds to cardiolipin in human cell lines

Cardiac muscle cells exhibit a higher density of mitochondria to meet their energy demands, rendering cardiomyocytes vulnerable to mitochondrial dysfunction and making them likely targets for mitochondrial toxic drugs.^34^ Previously, caspofungin-induced cardiomyopathy has been shown to impact patients receiving treatment for candidiasis, with mitochondrial dysfunction identified as a contributing factor^35,36^. Therefore, we investigated the effects of the caspofungin interaction with mitochondria in human heart cells using BD-CSF. Human hybrid cardiomyocyte cell line (AC16) and human embryonic kidney cells (HEK293) were treated with 15 µM BD-CSF followed by staining with MitoTracker Green or NAO. Following the co-staining, cells were analyzed using fluorescence microscopy. Our data demonstrate the co-localization of BD-CSF with both MitoTracker and the cardiolipin specific NAO stain suggesting that the CSF interaction with mitochondria occurs through cardiolipin association in human cells (Fig. 6A). Additionally, results from the spectral characterization of cells indicated an increased caspofungin uptake in cardiac cells relative to kidney cells which is consistent with the higher concentrations of mitochondria and cardiolipin present in cardiac cells (Fig. 6B). A detailed analysis of the spectral data (Supplementary Fig. 7) shows no statistical difference in fluorescence intensity of the markers (NAO or MitoTracker) between cells treated with CSF probes and those without irrespective of cell type. However, cells that were treated with BD-CSF prior to mitochondrial specific dyes, did not stain equally effectively with mitochondrial markers indicating that BD-CSF occupies the binding sites specific to either MitoTracker or NAO. These data strongly suggest that fluorescently labelled caspofungin binds to human mitochondria via its affinity for cardiolipin, in a manner analogous to its interaction with Cryptococcus cells.

**Figure 6.**
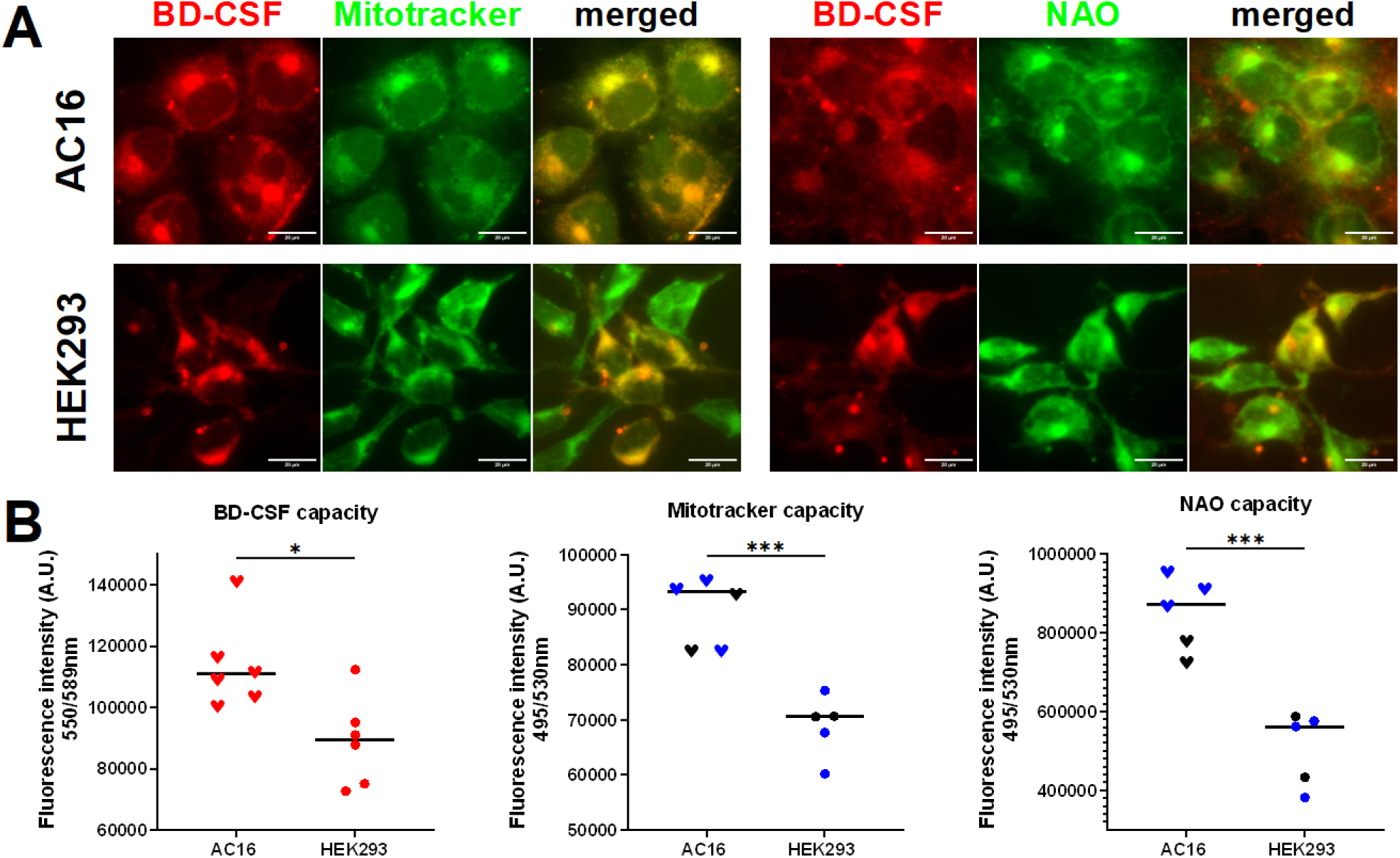
**A:** Representative images showing the co-localization of mitochondrial markers (NAO and MitoTracker) with (top panels) BD-CSF in the human heart (AC16) and (bottom panels) kidney (HEK293) cell lines. **B:** Increased capacity of AC16 cells for BD-CSF (red, left), MitoTracker (middle) and NAO (right) regardless of pre-treatment with unlabelled CSF (blue).

## Discussion

The existing ß-1,3-glucan synthase-targeting antifungals, such as caspofungin, are shown to inhibit glucan synthase activity by potentially binding to the extracellular surface of the plasma membrane bound Fks1 subunit. However, the detailed mechanisms by which these drugs perturb glucan synthase function in cells are not well understood and most of the available information on glucan synthase inhibition is based on the data obtained for *C. albicans* and *S. cerevisiae*^37-40^. *C. neoformans* is highly resistant to caspofungin treatment^6^ despite the *FKS1* gene being essential and the high sensitivity of glucan synthase to caspofungin *in vitro*^8^. To investigate the molecular basis of caspofungin resistance in *C. neoformans*, we first sought to determine whether the drug enters the yeast cells and, if so, to define its subcellular distribution relative to the glucan synthase complex. For this we designed and generated two different fluorescent derivatives of caspofungin – BD-CSF and F-CSF. We used these two chemically distinct fluorescent derivatives of caspofungin to ensure that the observed phenotypic changes in fluorescence uptake are related to the CSF moiety of the probe, rather than a fluorescence marker itself. We observed, using fluorescent microscopy, cytoplasmic signals for both fluorescent caspofungin probes within the wild-type *C. neoformans* instead of the expected plasma membrane binding. We performed a phenotypic characterization of the wild-type *C. neoformans* cells revealing phenotypic features present in all samples regardless of which fluorescent caspofungin derivative was used, thereby indicating that both probes localize in the same manner. Moreover, the results of the MIC assays were consistent with the known cryptococcal resistance to caspofungin showing that the apparent MIC is much higher than the clinically relevant CSF concentration of 10 µM. Notably, both BD-CSF and F-CSF exhibit MICs comparable to those of caspofungin. In this species, the primary restriction may be drug delivery rather than intrinsic potency; thus, even a slight reduction in binding affinity may not lead to a significant alteration in MIC. To determine if caspofungin is localized to the plasma membrane as predicted, we performed a co-localization study between our fluorescent BD-CSF probe and the ergosterol binding molecule, Filipin III, and found minimal signal overlap between the two probes indicating that the majority of fluorescent caspofungin is localized elsewhere.

Both fluorescent CSF probes were localized in the cytoplasm in a manner that resembled mitochondrial staining, leading us to characterize cryptococcal cells subjected to caspofungin treatment with the mitochondrial marker - MitoTracker. We observed significant overlap of MitoTracker and the fluorescent caspofungin derivatives, indicating that the drug localizes to and interacts with mitochondria in *C. neoformans*. Although this finding was surprising it is not unprecedented that mitochondria are targeted by drugs. Indeed, antifungal drugs, including echinocandins and azoles, induce a conserved oxidative damage pathway involving mitochondria-derived reactive oxygen species^41^. This oxidative stress, when not lethal, appears to trigger protective mitochondrial responses that facilitate survival. Persister populations in *C. albicans* are highly tolerant to fluconazole and mitochondrial oxidative stress response pathways are central to their survival during treatment^42^. The role of mitochondria in mediating not only tolerance but also virulence, suggests that mitochondrial homeostasis supports fungal fitness under antifungal stress in both *C. albicans* and *C. neoformans^43^*. The flexibility of Candida’s mitochondrial respiratory pathways was shown to affect mitochondrial function and enable mitigation of azole-induced stress effects^44^.

The apparent correlation between the MitoTracker and BD-CSF fluorescence was further confirmed by our colocalization study revealing a high Pearson coefficient R characteristic of the colocalization between the mitochondria and fluorescent caspofungin. We performed high-resolution microscopy to further characterize the apparent fluorescent caspofungin binding to mitochondria. We obtained images showing that BD-CSF/F-CSF fluorescence shares a phenotype and overlaps with MitoTracker, regardless of which fluorescent caspofungin was used, further confirming the fluorescent drug interaction with mitochondria.

Our attempts to show colocalization of BD-CSF fluorescence with NAO - the marker for the mitochondrial lipid, cardiolipin- was not successful due to the marker outcompeting the drug from mitochondria. Based on our imaging data showing BD-CSF/NAO competition we hypothesized that cardiolipin might be binding to the fluorescent caspofungin. We verified the direct interaction between cardiolipin and caspofungin using isothermal titration calorimetry. The average *K*_d_ values obtained from these experiments indicate strong binding of caspofungin to cardiolipin, comparable to specific NAO – cardiolipin binding. The average N-value of our caspofungin-cardiolipin ITC experiments suggests a ∼1.5:1 binding ratio of these molecules, similar to the 2:1 stoichiometry observed for the NAO-cardiolipin interaction. We did not observe reproducible binding of caspofungin to phosphatidylcholine (PC), the most abundant phospholipid in the mitochondrial membrane, further supporting the idea that caspofungin binds preferentially and likely only to cardiolipin.

Based on these data, we propose a model for caspofungin resistance in *C. neoformans* whereby the high affinity interaction of caspofungin with cardiolipin in the mitochondrial membrane sequesters the drug from the plasma membrane, lowering the effective drug concentration near glucan synthase, the intended target. In this model, the mitochondrion essentially acts as a “sponge” soaking up the introduced drug, preventing it from acting effectively as an antifungal. To test this model, we assumed that once the sponge is pre-saturated, e.g., with the cardiolipin specific compound NAO, caspofungin will regain its efficacy and the drug will be effective against *cryptococcal* infections at clinically relevant doses. Consistent with the model, results obtained for the cells pre-treated with NAO at the very early logarithmic stage of growth show almost 90-fold increase in the efficacy of caspofungin against NAO pre-treated cells compared to untreated cells. In parallel we performed the same set of experiments using BD-CSF. For the NAO pretreated samples, the efficacy of BD-CSF treatment was lower than for the parent drug (31-fold).

Additionally, we characterized the competition between caspofungin (CSF/ BD-CSF) and NAO for cardiolipin binding revealing that the derivatization of caspofungin increases the affinity of BD-CSF for cardiolipin. This increased affinity for cardiolipin explains the lowered efficacy of the BD-CSF as compared to caspofungin in the combinatory treatment. According to our data, BD-CSF has a cardiolipin-binding affinity that is more comparable to NAO than caspofungin, and thus outcompetes NAO from the pre-saturated mitochondria. As a result, the BD-CSF localization pattern is shifted away from the plasma membrane and Fks1, lowering the effective concentration of the drug despite the NAO pre-treatment intended to block the mitochondrial sponge.

We carefully considered the effect that the cryptococcal cell cycle might have on the observed drug response. In the experimental conditions used initially, we had a mixed cell population of budding and single cells out of which some were likely quiescent based on the length of the available culture growth data and the length of performed experiments. Based on the obtained data, we noticed that budding cells appear to be more affected by caspofungin treatment. Interestingly, for *C. albicans* it has been established that echinocandins are more effective against dividing yeast cells than against quiescent cells^16^. Moreover, it was previously shown that mitochondrial modulation drives age-associated fluconazole tolerance in *C. neoformans*^33^. Additionally, glucan synthase in *S. cerevisiae* appears to be most active enzymatically at early logarithmic growth stages^39^ and Fks1 in *C. neoformans* is internalized and removed from the plasma membrane in older cells^9^. Thus, for experiments testing the proposed mitochondrial “sponge” model, we initiated the experiments with cells cultured to the very early logarithmic stage. Significant differences in the size of budding, but not single, cells were observed upon exposure to NAO +/-caspofungin with the most dramatic decrease in the cell size for the cells exposed to both NAO and consecutive caspofungin treatment. The effect of BD-CSF on the size of budding cells was comparable to the results obtained with caspofungin, and additionally a statistically significant decrease was observed for cells pre-incubated with NAO and treated with BD-CSF. These combined observations lead to the conclusion that, as in *C. albicans,* caspofungin has the greatest effect on the population of budding cells.

Since, cryptococcal cells take up caspofungin and localize it to the mitochondria we also looked at the effects of caspofungin treatment on human cardiac and kidney cells. Interestingly, the cytotoxic effect of higher caspofungin concentrations on mitochondria in human cardiac cells has been observed^36,45^. These observations suggest that caspofungin binding to cardiolipin may be the cause of its cytotoxicity, preventing the use of increased drug concentrations to overcome the pathogen’s resistance. We found comparatively higher uptake of fluorescent caspofungin into heart cells as compared with the kidney cells, consistent with the higher volume of mitochondria as well as a role for cardiolipin in heart tissue. Our results suggest that the cardiotoxicity of caspofungin is caused by its binding to cardiolipin in cardiac mitochondria. These observations suggest that chemical modifications to caspofungin that reduce its binding to cardiolipin could increase its efficacy against Cryptococcus while reducing its cardiotoxicity.

Results presented here point to a previously unexplored phenomena related to the role of mitochondrial stress pathways in *C. neoformans* resistance to caspofungin. *C. neoformans* survives exposure to various stressors during growth in the human host, including oxidative stress, and it has been established that mitochondrial function plays a significant role in oxidative stress resistance^46,47^. Additionally, mitochondrial morphology has been shown to have significant impacts on both stress resistance and virulence^24^. According to our model, cryptococcal cells evade inhibition of glucan synthase by internalizing the drug and sequestering it by binding to the mitochondrial phospholipid cardiolipin. The capacity of this “sponge” effect is related to the number of mitochondria, their metabolic status, and the concentration of cardiolipin. It is likely that the reduced cell viability observed at higher caspofungin concentrations arises from its cytotoxic effects on mitochondria rather than from inhibition of Fks1.

The hydrophobic lipid side chain of pneumocandin B_0_, caspofungin precursor, has long been assumed to function as a membrane-anchoring “claw” that promotes productive engagement of the membrane-embedded glucan synthase Fks1^29,48^. Our data challenge this paradigm in *C. neoformans*. We show that the lipid tail displays higher affinity for mitochondrial cardiolipin than for the plasma membrane–associated target, resulting in sequestration of the drug away from Fks1 and functional loss of antifungal activity. Thus, in Cryptococcus, the very structural feature thought to confer potency instead drives off-target mitochondrial partitioning and therapeutic failure. These findings establish cardiolipin binding as a previously unrecognized determinant of echinocandin inefficacy and highlight lipid-tail chemistry as a critical, actionable parameter for next-generation echinocandin design. Future chemical and semisynthetic strategies should therefore prioritize decoupling membrane targeting from cardiolipin affinity to enable development of safer and more effective antifungals with improved pharmacokinetic and pharmacodynamic profiles.

## MATERIALS AND METHODS

### Fungal strains and media

*C. neoformans* strain KN99α was used as the wild-type strain. *C. albicans* WO1 strain was used as a control. Fungal strains were grown in YNB 2% (0.67% yeast nitrogen base, 2% dextrose, pH 7.0, with 50 mM MOPS [morpholinepropanesulfonic acid])

### Caspofungin derivatization

Detailed synthetic protocol is described in Supplementary Information for both BD-CSF and F-CSF derivatives.

Briefly, the BD-CSF derivative of caspofungin (Biosynth International, Inc. cat# FC16269) was produced by incubating BDP® 576/589-succinimidate (Lumiprobe, cat# 2G420) with pure caspofungin in the presence of triethylamine as a proton acceptor in DMF. BDP® 576/589 – NHS ester an activated N-hydroxysuccinimide ester of the dye with reactivity to primary and secondary amino groups.

The F-CSF derivative of caspofungin was synthesized in Fridman Lab according to the published synthetic protocol^17^.

The BODIPY-CSF derivative of caspofungin was gifted to M.D. by Chris Arnatt, Ph.D., Department of Chemistry, Department of Pharmacology and Physiology, Saint Louis University and was synthesized according to the published protocol^11,20^.

### MIC of caspofungin

Antifungal assays were performed according to the CLSI M27A method, and the amount of growth after incubation was compared to no-drug controls. MICs for each organism were determined by testing serial dilutions of caspofungin and its fluorescent derivatives in a 96-well microtiter broth assay. Cells were inoculated into YNB pH=7 medium to a final concentration of 4 × 10^3^ cells/ml *(C. albicans*) or 2× 10^3^ cells/ml (*C. neoformans*), incubated for 24 h (48 h for *C. neoformans*) at RT, followed by the measurement of OD_600_. The MIC was defined as the lowest concentration of drug that showed absence of growth compared to no-drug control.

### Fluorescence microscopy

Cells were imaged using an Olympus BX61 with a 100× objective or, where indicated in the text, Zeiss ELRYA7 high resolution microscope located in the Duke Light Microscopy Core. Images were processed using FiJI software^49^.

### Preparation of samples for microscopy characterization

Initially, we characterized KN99 cells cultured to mid logarithmic stage (OD_600_ 1-2) in YNB, pH= 7 media followed by culturing for additional 24h with/without 20-25 µM of CSF, BD-CSF, or F-CSF in the 30°C incubator with 300rpm shaking. Then cells were briefly (15-30min) stained with 1mg/mL solution of Calcofluor white (CFW; Sigma cat#F3543). Cell survival was determined by CFU from the aliquot of cells prior to CFW exposure.

For Filipin III staining, KN99 cells, cultured to OD_600_=4 in YNB media at pH 7, diluted to OD_600_=1 prior to 24-hour treatment with 10µM Caspofungin or BD-CSF probe. Concentration of the caspofungin was dropped to ensure that the drug won’t affect viability of the analyzed cells. Next, 5 µL of a 10 µg/mL Filipin III (Biogems cat#4804999) stock solution was added to the samples for a duration of up to 10 minutes, followed by a brief wash in PBS and immediate image collection to prevent artifacts associated with rapid oxidation of Filipin III.

Phenotypic characterization of cryptococcal mitochondria was performed using mid logarithmic (OD_600_ =1-4) KN99 cells, resuspended at OD_600_=1 prior to 24h treatment +/- 25 µM of CSF, BD-CSF, or F-CSF in the 30°C incubator with 300rpm shaking followed by staining with the carbocyanine-based dye MitoTracker Green FM (Invitrogen cat#M7514) or MitoTracker Deep Red FM (Thermo Fisher cat#M22426). MitoTracker staining was performed according to manufacturer’s recommendations. MitoTracker Green FM was used only for the initial phenotypic characterization of cryptococcal mitochondria following the CSF treatment. MitoTracker Deep Red FM was used in all of the subsequent experiments.

Nonyl Acridine Orange (NAO; Acridine Orange 10-Nonyl Bromide; Invitrogen cat#A1372) staining was performed with wild-type *C. neoformans* cells, cultivated to OD_600_ =1 in YNB (pH 7), followed by treatment with 10 µM BD-CSF for a duration of 18 hours. Following drug incubation and PBS washes, cells were stained with NAO by adding 0.5 µL of a 2 mM NAO stock per mL of culture (resulting in a final concentration of 1 µM, at 1×10⁸ cells/mL) and incubated for 30 minutes at 30 °C with shaking at 300 rpm.

### BD-CSF vs Caspofungin competition assay

KN99 cells, cultured to OD_600_=4 in YNB media at pH 7, diluted to OD_600_=1 were treated with 25 µM Caspofungin or BD-CSF for 90 minutes followed by extensive wash in DPBS and resuspension in the fresh media supplemented with the same or the opposite caspofungin probe for additional 90 minutes. The cells were washed prior to the collection of spectral data using SpectraMax iD3 (Molecular Devices) plate reader.

### Statistical analysis of cells

To quantify the prevalence of the fluorescent CSF derived phenotypes, three images taken from spatially distinct regions of the slide were analyzed to ensure that a representative cell population was assessed. For the statistical analysis of individual cells, 100 or more cells were identified by cellular perimeter in BF channel followed by reading of the fluorescence within that area from the reminder of channels. Using that approach we obtained datasets containing cell size (µm^2^) as well as the average fluorescence intensity of individual cells. The budding status of each cell was assessed using bright field (BF) images. Cells were classified as either “budding” or “single” in cases where no budding was observed. Mother and daughter cells were individually characterized, systematically accounted for, and quantified together under the “budding” category.

Colocalization analysis was done using Fiji software (plugin Coloc 2). It implements and performs the pixel intensity correlation over space methods of Pearson, Manders, Costes, Li and more, for scatterplots, analysis, automatic thresholding and statistical significance testing. Pearson’s coefficient for each cell was calculated for the fluorescent channels of interest.

Statistical analysis was done using Graph Prism software. Statistical significance of the observed differences was calculated using one-way ANOVA. One asterisk (*) identifies adjusted P values between 0.01 and 0.05, two asterisks (**) identify adjusted P values between 0.01 and 0.001, etc.

### Isothermal titration calorimetry

18:1 Cardiolipin (1’,3’-bis[1,2-dioleoyl-sn-glycero-3-phospho]-glycerol (cat# 710335)) and 18:1 (Δ9-Cis) PC (1,2-dioleoyl-sn-glycero-3-phosphocholine (cat# 850375)) were purchased from Avanti Research.

Isothermal titration calorimetry (ITC) experiments were performed using a VP-ITC microcalorimeter (Malvern Panalytical). All experiments were conducted in triplicates under atmospheric pressure at 25°C in a matching ITC buffer (EtOH/H2O=1/1) while stirring at 300 rpm. Cardiolipin was solubilized with the ITC buffer to a concentration of 1.0 mM and caspofungin to a concentration of 25 μM. All solutions were then stored o/n until use in 4°C followed by degassing for 30 minutes under vacuum immediately prior to the experiment. As a consequence of degassing, small differences in EtOH concentrations occur between the experimental repeats, resulting in the difficulties in precise determination of the concentrations of caspofungin and cardiolipin warrant cautious interpretation of the fit stoichiometry. Aliquots from a 0.5 mL syringe containing 1.0 mM CL were sequentially injected into a 2.50 mL sample cell containing a 25.0 μM solution of caspofungin. All titrations were performed using an initial injection of 2 μL over 4 seconds followed by 23 identical injections of 12 μL with a duration of 24 seconds per injection followed by an equilibration time of 450 seconds between injections (7.5 minutes). Control experiments were conducted with phosphatidyl choline (PC; 1.0 mM, EtOH/H2O=1/1) titrated into caspofungin (25 μM) using the same experimental parameters as described above. Appropriate blank data subtraction and data analysis were executed using the program ORIGIN 7 SR4 (v7.0552). Manual baseline corrections were applied to correct for heat of dilution background signal (possibly arising from small differences in EtOH concentrations in the sample cell and syringe).

### Human cell lines

AC16 and HEK293 cell lines were kind gift from Dr. Lefkowitz lab, Duke University, Durham, NC. AC16 and HEK293 cell lines were plated in Ti75 flasks in DMEM/F-12, HEPES, no phenol red media (GIBCO #11039021) supplemented with, pen strep and 10% FBS.

For the fluorescence intensity measurements, AC16 cells or H9C2 cells were seeded at a density of 5 × 10^3^ cells/well in 96-well poly-L Lysine coated plates and incubated at 37 °C in the humidified CO2 incubator. DPBS was added to the unused wells to prevent evaporation. After 24 h incubation for cell adhesion, the DMEM/F-12 medium (1% FBS) containing vehicle or drug was added to wells. Treated cells were then incubated for 2h at 37 °C in the humidified CO_2_ incubator followed by incubation with the DMEM/F-12 medium (1% FBS) containing MitoTracker Green or NAO respectively. Following wash with DMEM/F-12 medium, the fluorescence was determined using a plate reader.

Fluorescent images were taken from the cells treated like previously, but seeded onto the chambered cell culture slides (MatTek #CCS-8).

## Acknowledgements

The National Institute of Allergy and Infectious Diseases of the National Institutes of Health provided funding to M.J.D. and J.K.L. under grant number R01AI123407 and R35GM130290 to M.A.S. National Institutes of Health provided funding to Duke Light Microscopy Core under grant number NIH S10: 1S10OD28703-01. This material is also based upon work supported by the National Science Foundation Graduate Research Fellowship Program under Grant No. DGE 2139754 to P.K.E. Any opinions, findings, and conclusions or recommendations expressed in this material are those of the author(s) and do not necessarily reflect the views of the National Science Foundation.

