## Supplementary Information for "Mitochondrial cardiolipin sequestration of caspofungin underlies *Cryptococcus neoformans* inherent resistance and may contribute to cardiotoxicity"

**Caspofungin derivatization**

The **BD-CSF** derivative of caspofungin was synthesized in Duke Small Molecule Synthesis Facility according to the published synthetic protocol[1]

General chemistry methods and instrumentation

Low-resolution electrospray ionization mass spectra (ESI-MS) were acquired on an Agilent 6310 ion trap. Analytical RP-HPLC was performed on an Agilent 1200 LC instrument equipped with a diode array detector and an Eclipse Plus C18 reversed-phase column (3.5 µm, 4.6 x 150 mm). The flow rate was 1 mL/min. Solvent A was $H_2O$ (with 0.1% formic acid) and solvent B was $CH_3CN$.

Synthesis of BD-CSF

A solution of the NHS-ester (15 mg, 0.035 mmol) in DMF (300 uL) was added dropwise over 3-5 minutes to a dry-ice/acetonitrile cooled solution (~-50 ºC) of caspofungin (43 mg, 0.035 mmol) in DMF (800 uL). The reaction was allowed to warm to room temperature as the cooling bath melted. Stirring was continued overnight. The mixture was loaded directly onto a RediSep$R_f$, C18, 26 g column. A solvent gradient from 100% $H_2O$ (containing 0.1% formic acid) to 100% MeOH was run over 30 minutes. The fractions containing product were concentrated to dryness under reduced pressure. The resulting residue was triturated with $Et_2O$ (~10-15 mL) and insoluble material was removed at the vacuum. The filter cake was dried in vacuo giving the product as a free-flowing dark purple solid (7.0 mg, 14%, >95% purity). ESIMS: m/z = 1402.8 [$(M-H)^-$].

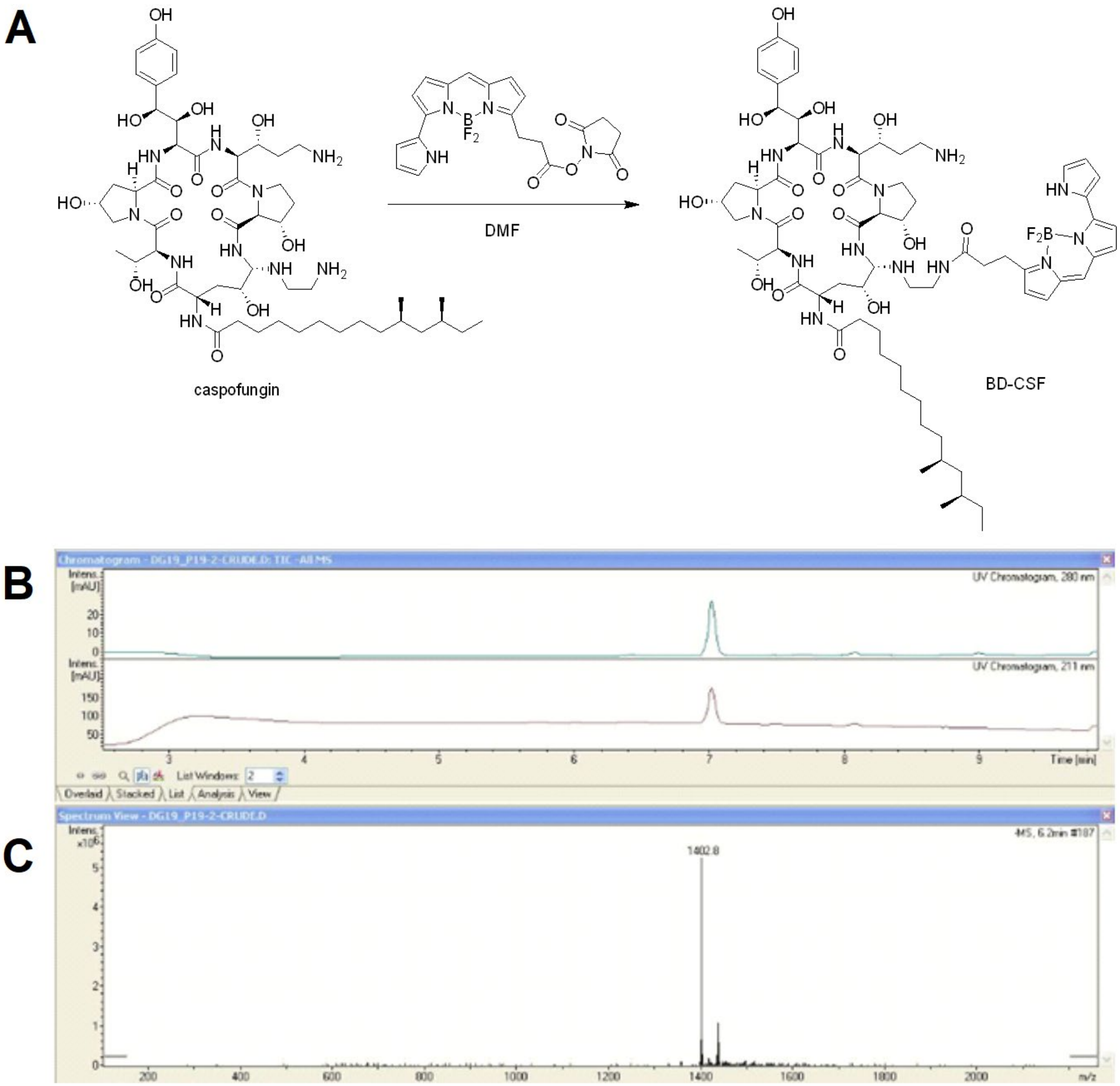


**Supplementary Figure 1 A:** The synthesis of BD-CSF. Procedure adapted from WO2015035102A2 (page 104) **B:** Analytic RP-HPLC chromatogram (diode array detector) of BD-CSF **C:** Electrospray ionization mass spectrum (ESI-MS) of BD-CSF consistent with the expected m/z = 1402.8 [(M-H)-].

The **F-CSF** derivative of caspofungin was synthesized in Fridman Lab according to the published synthetic protocol[2].

General chemistry methods and instrumentation

$^1$H-NMR spectra were recorded on BrukerAvance 400 MHz spectrometers. $^{13}$C-NMR spectra were recorded on BrukerAvance 400 or 500 MHz spectrometers at 100 MHz. Chemical shifts (reported in ppm) were calibrated to $CD_3OD$ ($^1$H: δ = 3.31, $^{13}$C: δ =49.0). Multiplicities are reported using the following abbreviations: s, singlet; d, doublet; t, triplet; dd, doublet of doublets; ddd, doublet of doublet of doublets; dt, doublet of triplets; m, multiplet. Coupling constants (*J*) are given in Hertz. High-resolution electrospray ionization (HRESI) mass spectra were measured on a Waters Synapt instrument. Low-resolution electrospray ionization mass spectra (ESI-MS) were measured on a Waters 3100 mass detector. Chemical reactions were monitored by thin-layer chromatography (TLC) (Merck, Silica gel 60 F254). Visualization was achieved using a cerium molybdate stain (5 g $(NH_4)_2Ce(NO_3)_6$, 120 g $(NH_4)_6Mo_7O_{24}\cdot 4H_2O$, 80 mL $H_2SO_4$, 720 mL $H_2O$) or with a UV lamp. All chemicals, unless otherwise stated, were obtained from commercial sources. Compounds were purified using Geduran Si 60 chromatography (Merck). The preparative reverse-phase high-pressure liquid chromatography (RP-HPLC) system used was an ECOM system equipped with a 5-µm, C-18 Phenomenex Luna Axia column (250 mm x 21.2 mm). Analytical RP-HPLC was performed on a VWR Hitachi instrument equipped with a diode array detector and an Alltech Apollo C18 reversed-phase column (5 m, 4.6 x 250 mm). The flow rate was 1 mL/min. Solvent A was 0.1% trifluoroacetic acid (TFA) in water, solvent B was acetonitrile. The SpectraMax i3x Platform spectrophotometer from Molecular Devices was used for fluorescence measurements.

Synthesis of F-CSF

Fluorescein was functionalized with the azide- according to previously reported procedures[3,4].

**CSF** was functionalized with a propargyl group on the phenol group according to previously reported procedure[2] . Then, compound **1a** (23.8 mg, 0.016 mmol, 1 eq) and azide-functionalized fluorescein (14.7 mg, 0.032 mmol, 2 eq) were dissolved in dry DMF (2 mL). A catalytic amount of $CuSO_4 \cdot 5H_2O$ and sodium ascorbate were added to the solution. The reaction solution was stirred at ambient temperature overnight and progress was monitored by ESI-MS, following the disappearance of **1a** ($[M-H]^-$ m/z 1430.6) and the appearance of **Boc-protected F-CSF** ($[M+H]^+$ m/z 1891.8). Upon completion of the reaction, the solvent was removed by lyophilization. The crude powder was then dissolved in isopropanol (1 mL), and 32% HCl (0.38 mL) was slowly added dropwise. The reaction was stirred at ambient temperature for 2 h. Progress was monitored by MS-ESI, following the disappearance of **Boc-protected F-CSF** ($[M+H]^+$ m/z 1891.8) and the appearance of **F-CSF** ($[M-H]^-$ m/z 1588.5). Upon completion, the solution was diluted with a solution of 80% acetonitrile 20% $H_2O$. Purification was done by preparative RP-HPLC (mobile phase: acetonitrile in $H_2O$ containing 0.1% TFA; gradient from 10% to 90%; flow rate: 20 mL/min) yielded the hydrochloride salt of **F-CSF** (15 mg, 54%) as an orange powder. HRESI-MS m/z calculated for $C_{79}H_{109}N_{14}O_{21}$, 1589.7892; found $[M+H]^+$, 1589.7889. $^1H$ NMR (400 MHz, $CD_3OD$) δ (ppm) 8.40 (d, *J* = 1.0 Hz, 1H), 8.18 (dd, *J* = 8.0, 1.6 Hz, 1H), 8.11 (s, 1H), 7.29 (d, J = 8.0, 1H), 7.23 (d, *J* = 8.7 Hz, 2H), 6.98 (d, *J* = 8.7 Hz, 2H), 6.69 (d, *J* = 2.3 Hz, 2H), 6.59 (d, *J* = 8.7 Hz, 2H), 6.53 (ddd, J = 8.7, 2.08, 2.04 2H), 5.14 (s, 2H), 5.00 (d, *J* = 3.2 Hz, 1H), 4.89-4.94 (m, 2H), 4.47-4.62 (m, 6H), 4.36 (d, *J* = 7.8 Hz, 1H), 4.28-4.33 (m, 2H), 4.19-4.24 (m, 2H), 4.02-4.13 (m, 2H), 3.97 (dd, *J* = 11.2, 3.1 Hz, 1H), 3.76-3.88 (m, 3H), 3.45-3.51 (m, 2H), 2.99-3.17 (m, 5H), 2.88-2.99 (m, 1H), 2.44 (dd, *J* = 13.0, 6.8 Hz, 1H), 2.20-2.32 (m, 5H), 1.94-2.16 (m, 5H), 1.77-1.89 (m, 1H), 1.52-1.65 (m, 2H), 1.20-1.51 (m, 15H), 1.17 (d, *J* = 6.1 Hz, 3H), 1.02 -1.10 (m, 2H), 0.81-0.92 (m, 10H). $^{13}C$ NMR (100 MHz, $CD_3OD$) δ (ppm) 176.7, 174.5, 173.7, 173.4, 172.8, 172.7, 170.5, 168.9, 168.5, 161.7, 159.6, 154.2, 145.0, 137.7, 135.5, 135.1, 130.2, 129.6, 128.7, 125.8,

125.6, 125.0, 116.0, 113.9, 110.9, 103.6, 77.4, 75.4, 75.0, 72.4, 71.3, 68.6, 68.3, 68.0, 65.3, 62.8, 62.5, 58.3, 57.1, 56.1, 56.0, 50.6, 47.1, 45.9, 43.0, 39.1, 38.5, 38.1, 37.8, 36.9, 35.6, 34.6, 32.9, 31.2, 31.1, 30.9, 30.8, 30.5, 30.3, 28.0, 27.1, 20.7, 20.2, 19.9, 11.6.

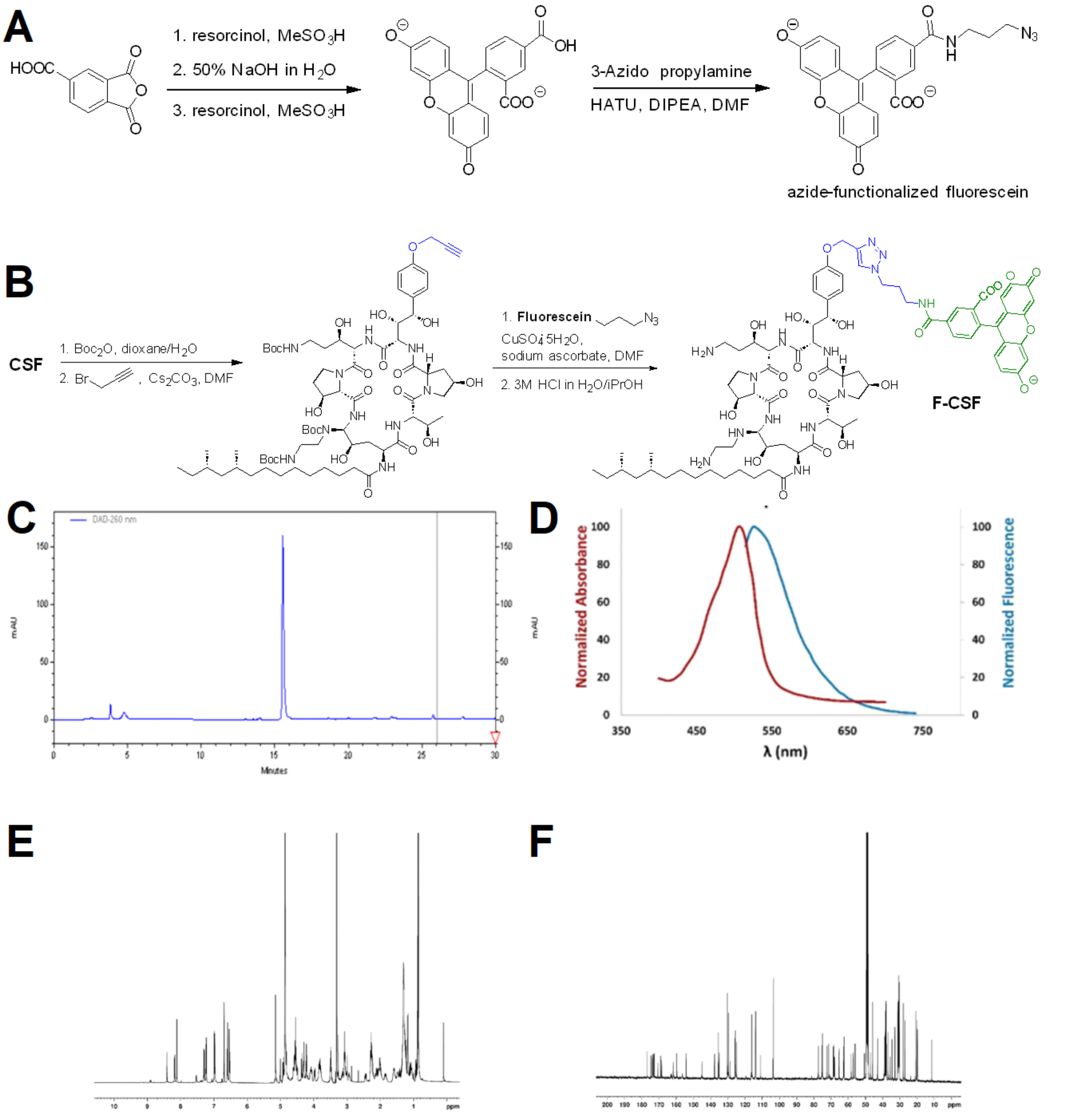


**Supplementary Figure 2 A:** The synthesis of the azide-functionalized fluorescein **B:** The synthesis of F-CSF. **C:** Analytic RP-HPLC chromatogram (diode array detector) of F-CSF **D:** Normalized absorption and emission spectra of fluorescein-labelled CSF. The measurements were made at the concentration of 10 µM in PBS (pH 7.4). **E:** 400 MHz $^{1}H$ NMR spectra of **F-CSF** in $CD_3OD$.**F:** 100 MHz $^{13}C$ NMR spectra of **F-CSF** in $CD_3OD$.

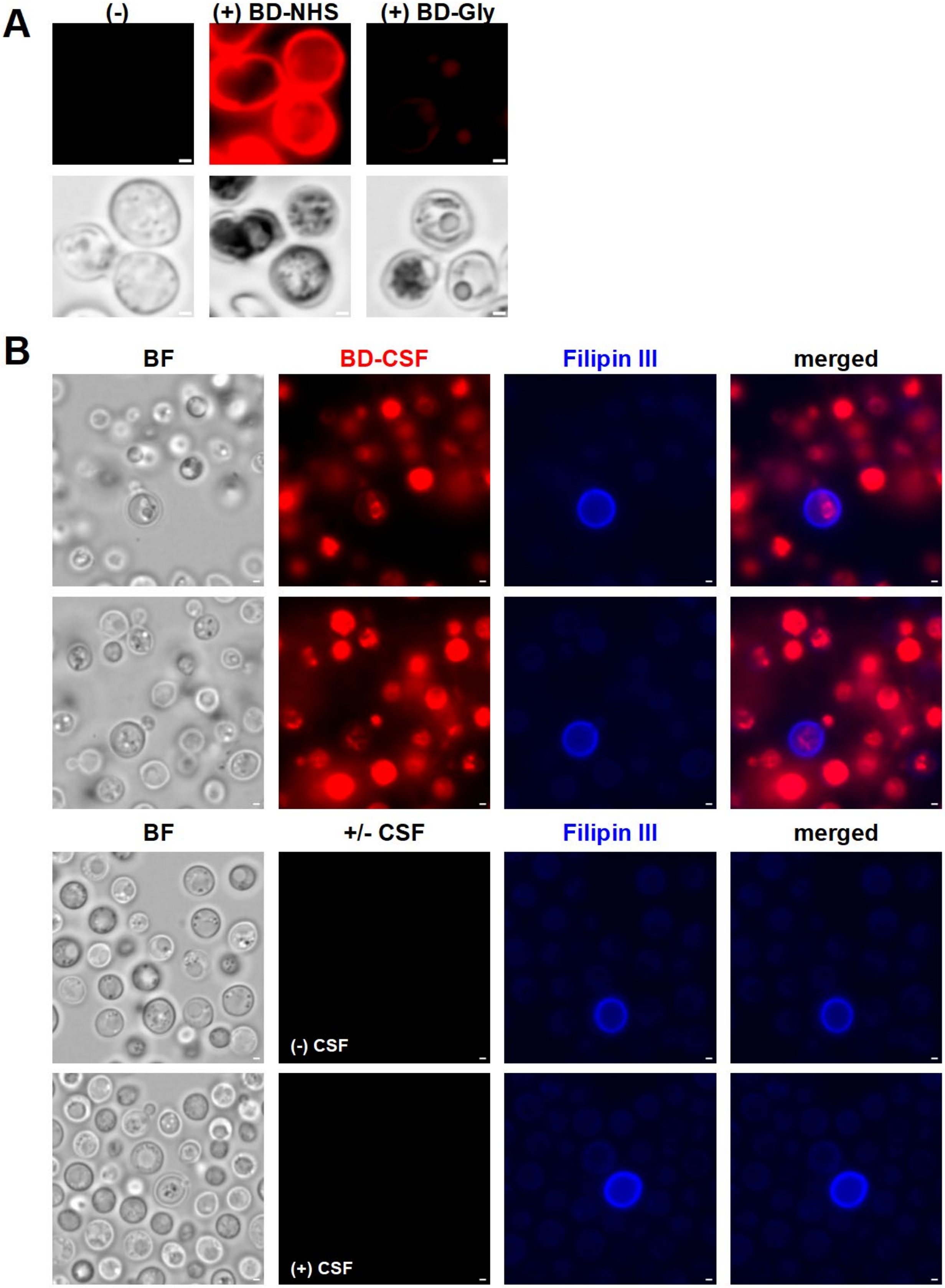
A
(-)
(+) BD-NHS
(+) BD-Gly
B
BF
BD-CSF
Filipin III
merged
BF
+/- CSF
Filipin III
merged
(-) CSF
(+) CSF

101

**Supplementary Figure 3 A:** Representative images of wild-type *C. neoformans* cells treated with 100µM BD-NHS and glycine derivatized with BD-NHS. **B:** Broad-field images of wild-type C. neoformans cells stained with Filipin III (blue) following the treatment with BD-CSF (red) or +/- CSF as a control. Scale bar within each image marks 1µm length.

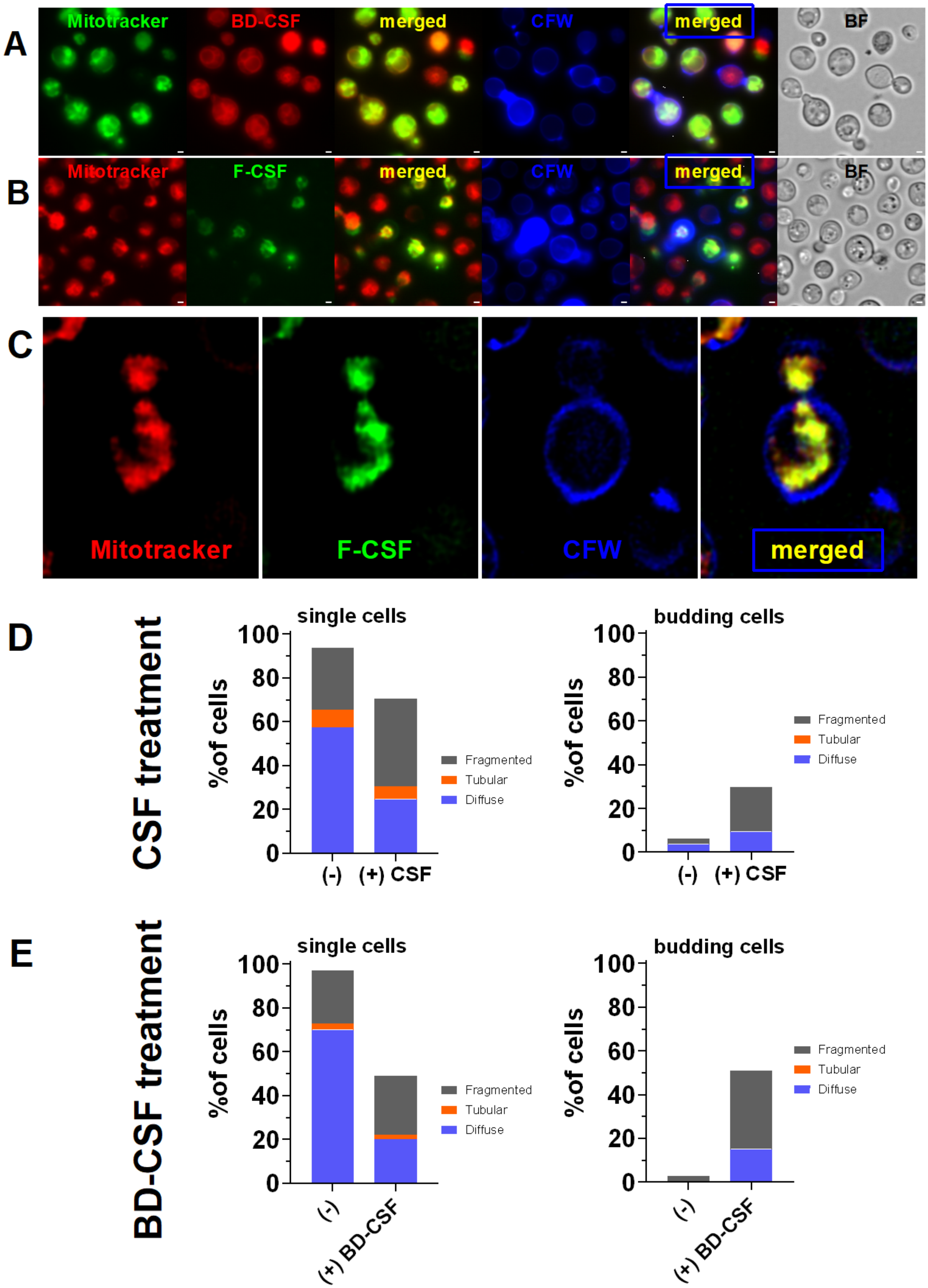
A
Mitotracker
BD-CSF
merged
CFW
merged
BF
B
Mitotracker
F-CSF
merged
CFW
merged
BF
C
Mitotracker
F-CSF
CFW
merged
D
CSF treatment
single cells
%of cells
100
80
60
40
20
0
(-)
(+) CSF
Fragmented
Tubular
Diffuse
budding cells
%of cells
100
80
60
40
20
0
(-)
(+) CSF
Fragmented
Tubular
Diffuse
E
BD-CSF treatment
single cells
%of cells
100
80
60
40
20
0
(-)
(+) BD-CSF
Fragmented
Tubular
Diffuse
budding cells
%of cells
100
80
60
40
20
0
(-)
(+) BD-CSF
Fragmented
Tubular
Diffuse

**Supplementary Figure 4 A:** Representative wide field images of wild-type *C. neoformans* cells treated with 20µM BD-CSF (red) co-labelled with MitoTracker (green) and calcofluor-white (blue) collected using Olympus microscope. Scale bar within each image marks 1µm length. **B:** Representative wide field images of wild-type *C. neoformans* cells treated with 20µM F-CSF (green) co-labelled with MitoTracker Deep Red FM (red) and calcofluor-white (blue) collected using Olympus microscope. Scale bar within each image marks 1µm length **C**: Representative images of wild-type *C. neoformans* cells treated with 20µM F-CSF (green) co-labelled with MitoTracker Deep Red FM (red) and calcofluor-white (blue) collected using the high-resolution Elyra 7 microscope. Phenotypic distribution of cells treated with **D:** CSF and **E:** BD-CSF.

**A**

**(1) CSF + CL** **(2) CSF + CL** **(3) CSF + CL**

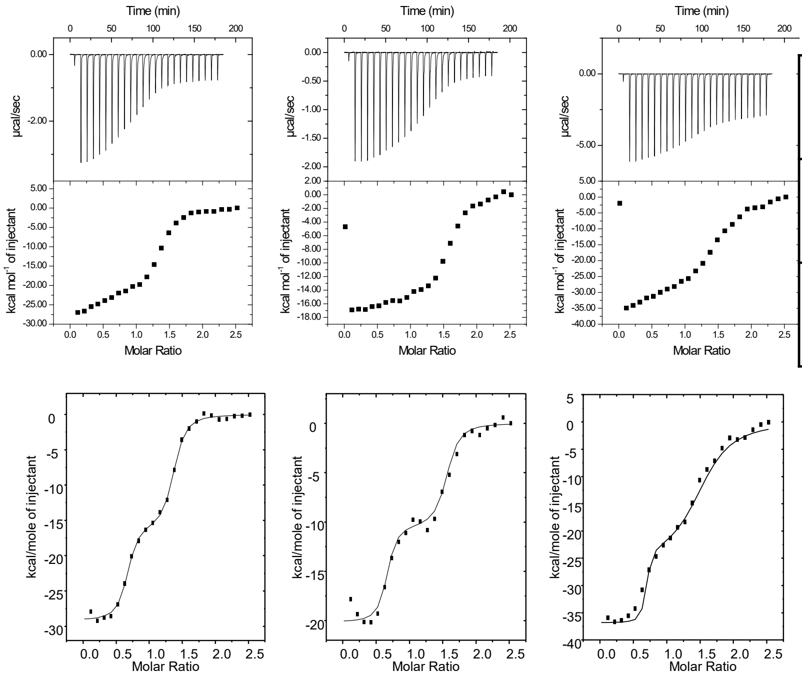


| | | |
|---|---|---|
| (1) CSF + CL | $K_{d(1)}$: $3.82 \times 10^{-9}$ M<br>$N_1$: 0.624 | $K_{d(2)}$: $2.62 \times 10^{-7}$ M<br>$N_2$: 0.707 |
| (2) CSF + CL | $K_{d(1)}$: $2.16 \times 10^{-9}$ M<br>$N_1$: 0.620 | $K_{d(2)}$: $2.30 \times 10^{-7}$ M<br>$N_2$: 0.903 |
| (3) CSF + CL | $K_{d(1)}$: $3.82 \times 10^{-9}$ M<br>$N_1$: 0.644 | $K_{d(2)}$: $1.72 \times 10^{-6}$ M<br>$N_2$: 0.874 |

**Average $K_d$:**

$K_{d(1)} = 3.26 \times 10^{-9} \pm 5.54\times 10^{-10}$ M

$K_{d(2)} = 7.39 \times 10^{-7} \pm 4.93 \times 10^{-7}$ M

**Caspofungin(CSF) binds strongly to Cardiolipin (CL)**

**B**

**(1) CSF + PC** **(2) CSF + PC** **(3) CSF + PC**

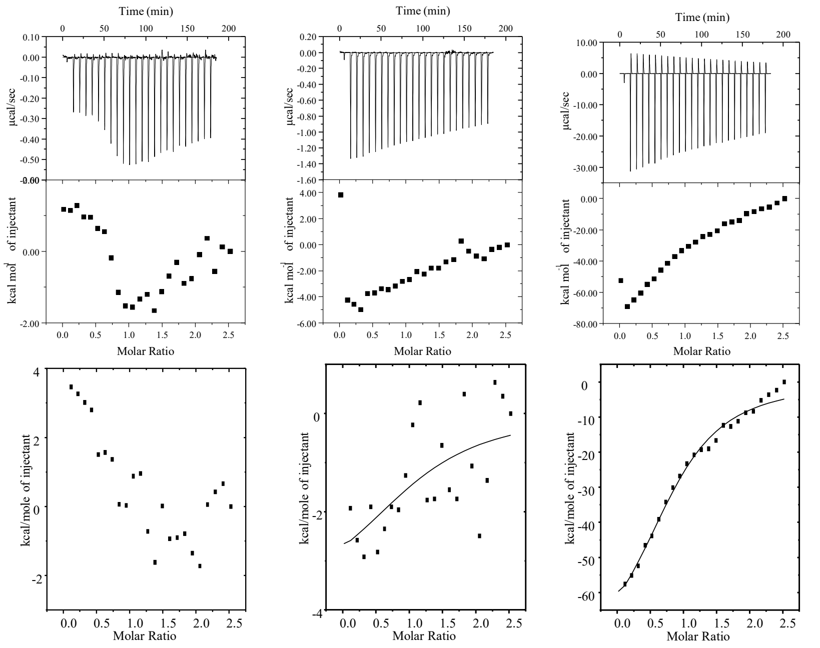


| | |
|---|---|
| (2) CSF + PC | $K_d = 1.36 \times 10^{-5}$ M<br>N: 1.2 |
| (3) CSF + PC | $K_d = 7.75 \times 10^{-6}$ M<br>N: 0.89 |

**Average $K_d$:**

$K_d = 1.08 \times 10^{-5} \pm 2.91\times 10^{-6}$ M

**Caspofungin (CSF) binds weakly and irreproducibly to phosphatidyl choline (PC).**

**Supplementary Figure 5** Isothermal titration calorimetry thermograms and resulting binding isotherms revealing two-site binding between **A:** caspofungin (CSF) and cardiolipin (CL) and one-site binding between **B:** caspofungin (CSF) and phosphocholine (PC). Experiments performed in triplicate. Dissociation constants ($K_d$) and stoichiometry values derived from binding isotherms are listed in their respective tables.

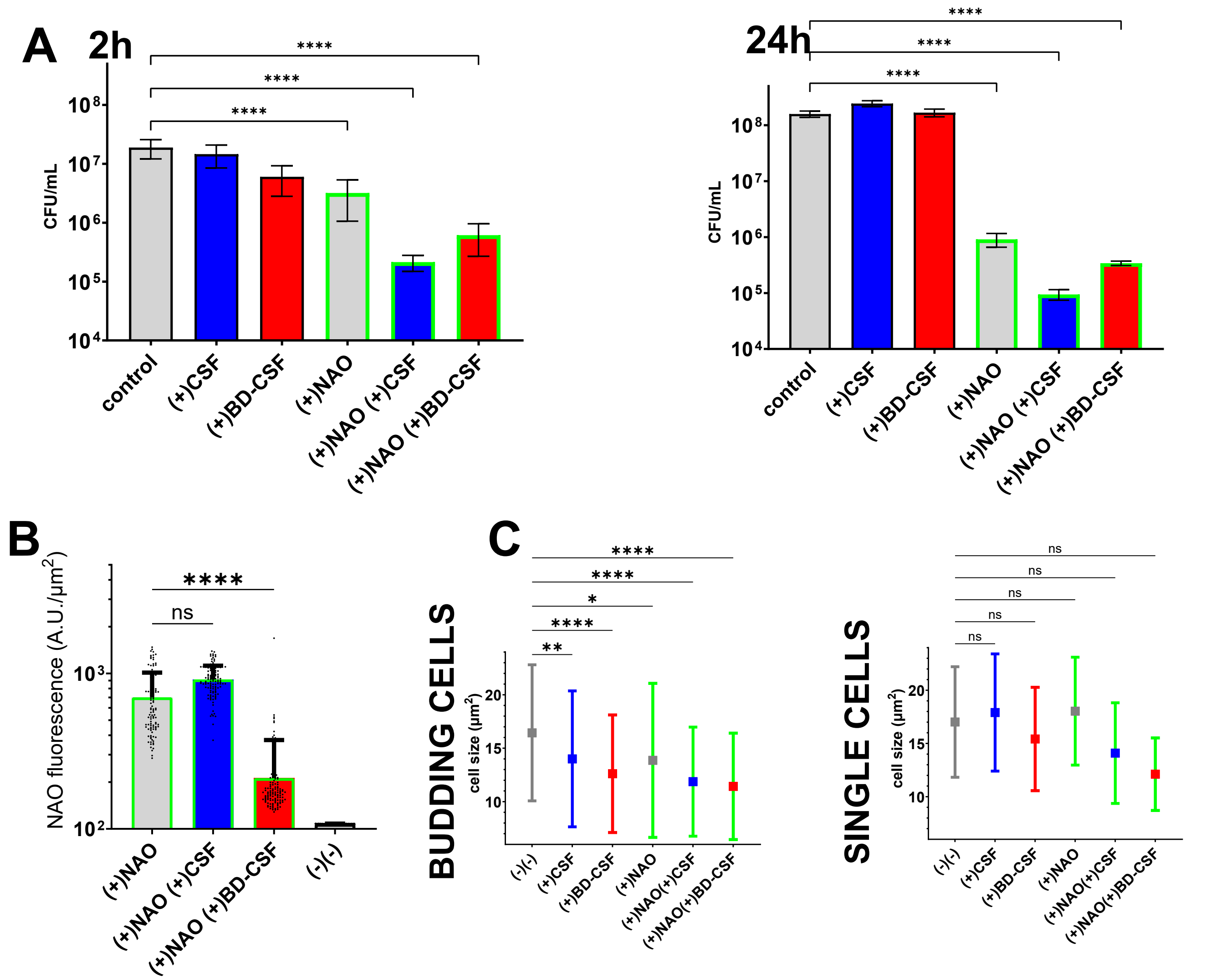


**Supplementary Figure 6 A:** Viability assay for wild-type *C.neoformans* (gray) treated with CSF (blue) or BD-CSF (red) with and without pre-treatment with 3μM NAO (green frame). Cells were treated with CSF/BD-CSF for2h and 24h following the 1h NAO pretreatment. **B:** Decrease of NAO fluorescence (green frame) following 10 μM BD-CSF treatment (red bar, green frame) as compared to no drug control (gray bar, green frame) and 10 μM CSF treatment (blue bar, green frame) **C:** Average cell size distribution of budding (left) and single (right) cells before (gray marker) and after (blue marker) CSF or (red marker) BD-CSF treatment preceded by NAO staining (green error bars). Statistical significance of the observed differences was calculated based on 100 or more representative cells per experimental condition, using one-way ANOVA. One asterisk (*) identifies adjusted P values between 0.01 and 0.05, two asterisks (**) identify adjusted P values between 0.01 and 0.001, etc.

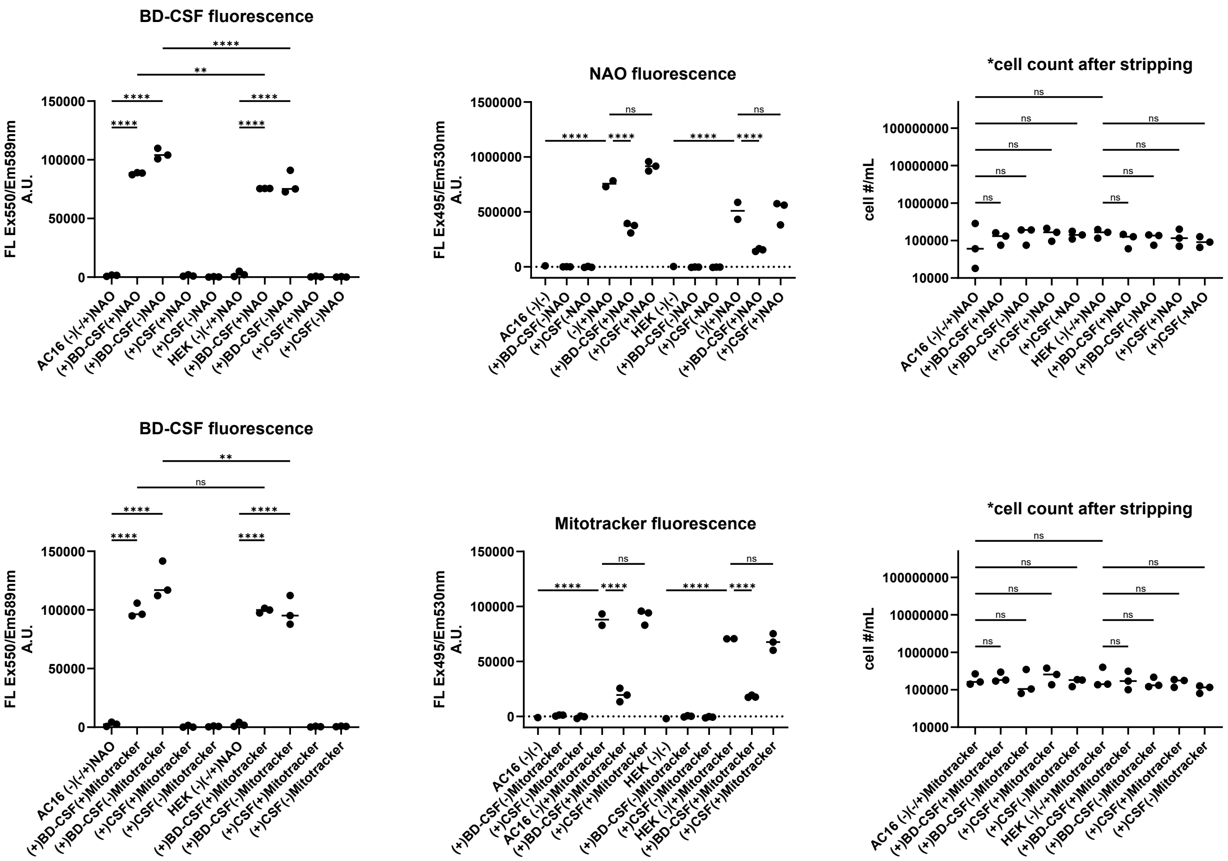


**Supplementary Figure 7** Statistical analysis of the fluorescence data obtained from cells treated with 15µM CSF/BD-CSF and then co-stained with NAO or MitoTracker. Individual wells were then stripped and the number of cells in each well was calculated.
